# A WDR35-dependent coatomer transports ciliary membrane proteins from the Golgi to the cilia

**DOI:** 10.1101/2020.12.22.423978

**Authors:** Tooba Quidwai, Emma A. Hall, Margaret A. Keighren, Weihua Leng, Petra Kiesel, Jonathan N. Wells, Laura C. Murphy, Joseph A. Marsh, Gaia Pigino, Pleasantine Mill

## Abstract

Intraflagellar transport (IFT) is a highly conserved mechanism for motor-driven transport of cargo within cilia, but how this cargo is selectively transported to cilia and across the diffusion barrier is unclear. WDR35/IFT121 is a component of the IFT-A complex best known for its role in ciliary retrograde transport. In the absence of WDR35, small mutant cilia form but fail to enrich in diverse classes of ciliary membrane proteins. In *Wdr35* mouse mutants, the IFT-A peripheral components are degraded and core components accumulate at the transition zone. We reveal deep sequence homology and structural similarity of WDR35 and other IFT-As to the coatomer COPI proteins *α* and *β*′, and demonstrate an accumulation of ‘coat-less’ vesicles which fail to fuse with *Wdr35* mutant cilia. Our data provides the first *in situ* evidence of a novel coatomer function for WDR35 likely with other IFT-A proteins in delivering ciliary membrane cargo from the Golgi necessary for cilia elongation.

## Introduction

The primary cilium is a highly-specialized sensory organelle and signaling hub compartmentalized from the rest of the cell and positioned with a unique interface towards the extracellular environment. Analogous to a cell’s antenna, many roles for cilia have emerged in development, disease and homeostasis ***(Reiter and Leroux, 2017)***. Whilst enrichment of signaling receptors and effectors in ciliary membranes is critical for cilia function, all biosynthesis of cilia-localised membrane proteins occurs in the endoplasmic reticulum and is sorted by the Golgi and efficiently routed via vesicles for incorporation into the elongating cilia membrane. The exact nature of this highly efficient transport process for the delivery of diverse cargos between membrane compartments remains unclear.

In mammalian cells, electron microscopy (EM) studies reveal the Golgi stacks closely apposed to the mother centriole ***(Sorokin, 1962; Wheatley, 1969)***. During intracellular ciliogenesis, small vesicles are observed to be recruited, most likely from the Golgi, to the mother centriole, which fuse to form a large preciliary vesicle (PCV) attached at the distal appendages ***(Yee and Reiter, 2015)***. More secondary vesicles then fuse with the PCV allowing elongation of cilia. Interestingly, the Golgi remains close to mature cilia, suggesting a continuous exchange of materials, enabling cilia maintenance ***(Sorokin, 1962; Wheatley, 1969)***. Several proteins essential for ciliogenesis localize to both the Golgi and the mother centriole and are implicated in this early stage of ciliogenesis including CCDC41 (CEP83), IFT20, HOOK2, and CEP164 ***(Baron Gaillard et al., 2011; Follit et al., 2006; Graser et al., 2007; Joo et al., 2013; Schmidt et al., 2012; Tanos et al., 2013)***. In some cases, including HOOK2 and CEP164, these components recruit Rab8a and Rabin-8, which facilitate membrane transport to cilia ***(Baron Gaillard et al., 2011; Moritz et al., 2001; Nachury et al., 2007)***. For some specific ciliary cargos, including rhodopsin ***(Wang and Deretic, 2014)*** and PKD2 ***(Follit et al., 2008, 2006; Hoffmeister et al., 2011; Kim et al., 2014; Noda et al., 2016)***, Golgi-to-cilia transport mechanisms have been described. However, these processes seem to involve cargo-specific trafficking modules. A more universal Golgi-to-cilia transport machinery, if one exists, has yet to be identified.

In contrast, movement of cargos within cilia requires highly conserved motor-driven macromolecular cargo binding complexes which traffic along axonemal microtubules closely apposed against the ciliary membrane, in a process known as intraflagellar transport (IFT) ***(Cole, 2009; Kozminski et al., 1993; Pazour et al., 1998; Pigino et al., 2009; Rogowski et al., 2013; Rosenbaum and Witman, 2002)***. Bidirectional movement of IFT complexes regulates cilia content; the IFT-B complex aids in kinesin-dependent anterograde transport of cargo, whilst the IFT-A complex helps in retrograde transport driven by dynein motors ***(Blacque et al., 2006; Efimenko et al., 2006; Jonassen et al., 2012; Lee et al., 2008; Piperno et al., 1998; Tran et al., 2008; Tsao and Gorovsky, 2008)***. The IFT-A complex is composed of three core (IFT144/WDR19, IFT140, IFT122/WDR10) and three peripheral proteins (IFT139/TTC21B/THM1, IFT121/WDR35, and IFT43) ***(Behal et al., 2012; Hirano et al., 2017; Piperno et al., 1998)***. However beyond classical retrograde defects (an inappropriate accumulation of cargos within the cilium) mutations in *IFT144, IFT140, IFT122, IFT121/WDR35*, and *IFT43* result in either severe reduction in cilia length or complete loss of cilia, implying they have critical roles in transport of cargo to cilia ***(Avidor-Reiss et al., 2004; Caparrós-Martín et al., 2015; Duran et al., 2017; Hirano et al., 2017; Liem et al., 2012; Mill et al., 2011; Takahara et al., 2018; Zhu et al., 2017)***. Indeed, several IFT-A mutants fail to localize a range of ciliary membrane proteins including EVC1/2, SMO, ARL13B, INPP5E, and SSTR3 to cilia ***(Caparrós-Martín et al., 2015; Fu et al., 2016; Jensen et al., 2010; Lee et al., 2008; Liem et al., 2012; Takahara et al., 2018)***. However, the mechanism of transport and the location of any IFT-A extra-ciliary function remains unclear.

The movement of cargos between membranes of spatially separated organelles in the cytoplasm involves vesicular trafficking. Indeed, IFT proteins have been observed to localize to various endomembranes and vesicular compartments outside cilia. For example, IFT20 localizes to the Golgi ***(Follit et al., 2006; Noda et al., 2016)***, whereas both IFT-B proteins (IFT20, IFT27, IFT46, IFT52, IFT57, IFT88 and IFT172) and IFT-A proteins (IFT139, IFT140) cluster around vesicles shown by immuno-EM and light microscopy ***(Sedmak and Wolfrum, 2010; Wood et al., 2012; Wood and Rosenbaum, 2014)***. Direct interaction of IFTs with membranes *in vitro* has also been described where the adaptor TULP3 and phosphoinositides mediate the membrane association of IFT-As ***(Mukhopadhyay et al., 2010)***. More recently, purified IFT172 was shown to bind to lipids and pinch off smaller vesicles, similar in size to classic COPI vesicles ***(Wang et al., 2018)***. It has been postulated that IFT proteins have evolved from coatomers: soluble multimeric protein complexes which ‘coat’ donor membranes, facilitating cargo enrichment and membrane remodeling prior to trafficking and fusion with target membranes ***(Jékely and Arendt, 2006; van Dam et al., 2013)***. Nonetheless, a functional requirement for an IFT-dependent Golgi-to-cilia trafficking module and what its dynamic architecture may resemble is currently unknown.

To dissect how trafficking of newly synthesized ciliary membrane proteins from the Golgi to the cilium occurs, we undertook a series of biochemical and imaging experiments in *Wdr35* null mouse embryonic fibroblasts (MEFs) ***(Caparrós-Martín et al., 2015; Mill et al., 2011)***. To distinguish extraciliary functions from canonical retrograde trafficking defects, we compared *Wdr35*^−/−^ phenotypes with those of the retrograde IFT dynein *Dync2h1*^−/−^ ***(Criswell et al., 1996; Huangfu and Anderson, 2005; Porter et al., 1999; Signor et al., 1999)***. Whilst retrograde accumulations of intact IFT-B were observed in both mutants, only in the absence of WDR35, the IFT-A holocomplex is fragmented and fails to enter the cilia in *Wdr35*^−/−^ cells. In addition to broad defects in the ciliary import of diverse membrane and lipidated proteins, we reveal an accumulation of ‘coat-less’ vesicles around the base of *Wdr35* mutants, which fail to fuse with the ciliary sheath. Together with our localization data, our results support the first *in situ* existence of a WDR35-dependent coatomer required to deliver essential cargo from the Golgi into cilia.

## Results

### Wdr35 null cells have rudimentary short cilia with intact transition zones but unstable axonemes

We utilized primary MEFs carrying null mutations in two components of the retrograde IFT machinery, the retrograde dynein heavy chain *Dync2h1* and the peripheral IFT-A component *Wdr35*, in order to unpick the stage at which ciliogenesis defects occurred ***(Caparrós-Martín et al., 2015; Mill et al., 2011)***. Cilia length measured by acetylated *α* tubulin staining was drastically reduced in both *Wdr35*^−/−^ (0.48µm mean ± 0.35 SD) and *Dync2h1*^−/−^ (0.76µm mean ± 0.35 SD) mutants compared to wild type (WT) (2µm mean ± 0.45 SD) MEFs (**Figure 1A, B**). Given there was no reduction in cilia number (**Figure 1C**), as previously shown ***(Fu et al., 2016; Liem et al., 2012; Mukhopadhyay et al., 2010)***, our results suggest that DYNC2H1 and WDR35 are needed for cilia elongation at later stages of ciliogenesis. Whilst defects in centriolar satellite trafficking, implicated in ciliogenesis, were previously reported for WDR35 mutant RPE-1 cells ***(Fu et al., 2016)***, we saw no difference in levels or localization of endogenously tagged PCM-1 protein (PCM1-SNAP) which marks centriolar satellites in *Wdr35*^−/−^ MEFs (**Figure 1 Supplement 1**). In *C. elegans* IFT-A mutants, defects in transition zone assembly gating were recently reported, where non-core IFT-A mutants allowed extension of the MKS module into the axoneme due to failure of cargo retrieval ***(Scheidel and Blacque, 2018)***. However, we observed intact transition zone modules as shown by NPHP1 and MKS1 localization in both mammalian mutants (**Figure 1D**). We noted that *Wdr35*^−/−^ axonemes, while acetylated, were not polyglutamylated suggesting differences in tubulin post-translational modifications (PTMs) and stability (**Figure 1D**). To bypass differences in tubulin PTMs between mutants, we compared the overall length of cilia by thick section TEM imaging of WT, *Dync2h1*^−/−^, and *Wdr35*^−/−^ MEFs. Short club-shaped cilia were observed in *Wdr35*^−/−^ (average 0.5µm) and *Dync2h1*^−/−^ (0.5µm-3µm) cells compared to normal 2-5µm long control cilia by TEM (**(Figure 1E**). Whilst the overall reduction in length of cilia as measured by TEM was consistent with acetylated *α* tubulin staining observed in *Wdr35*^−/−^ mutants, the acetylated axoneme was much shorter compared to the overall length measured by TEM in *Dync2h1*^−/−^ cells. Together this data suggests that the initial steps of ciliogenesis occur in both *Dync2h1*^−/−^ and *Wdr35*^−/−^ mutants, however, subsequent axoneme elongation is differentially affected.

**Figure 1.**
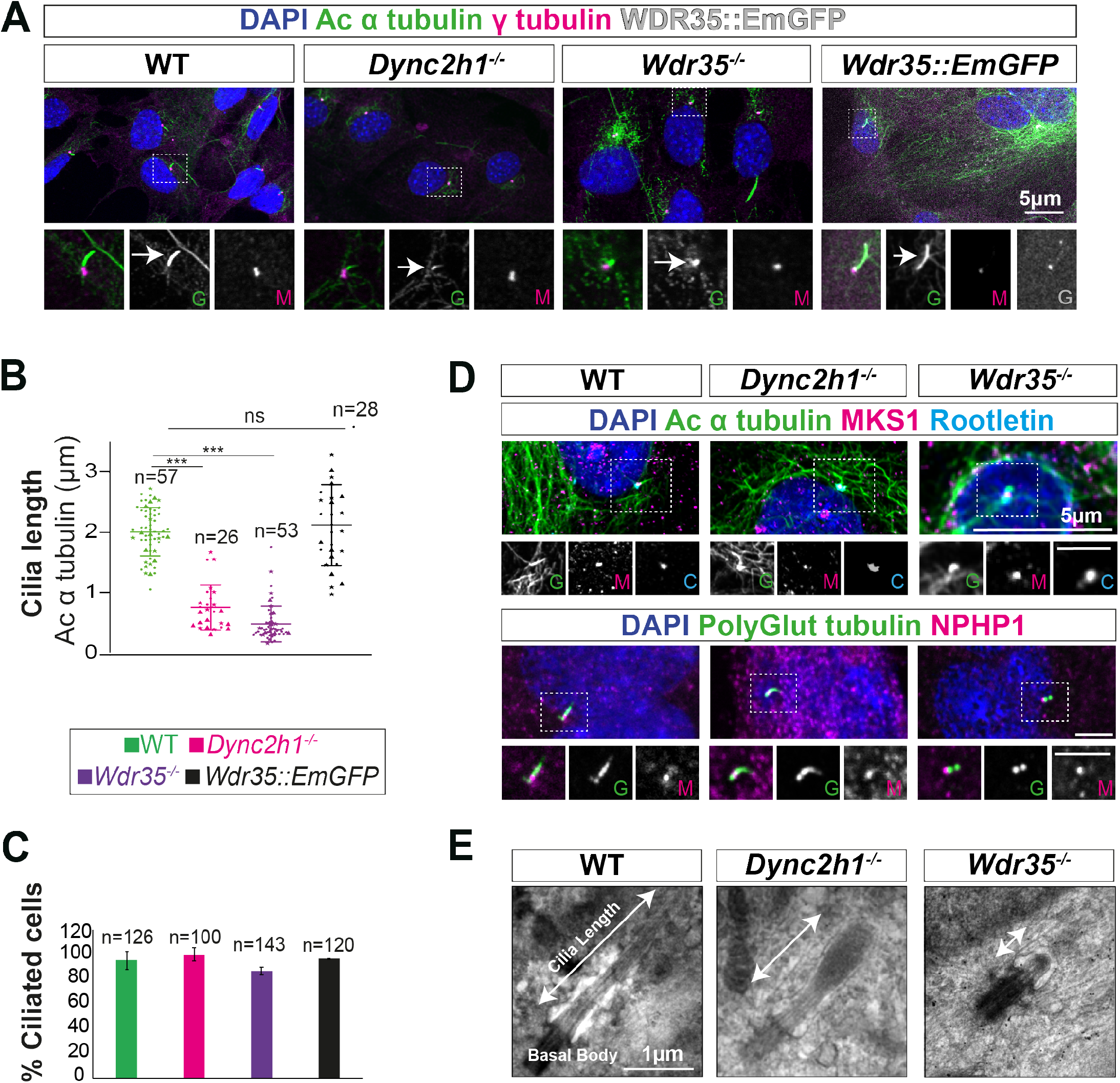
Wdr35^−/−^ and Dync2h1^−/−^ mutant cells have a drastic reduction in cilia length but have no difference in the number of cilia. (A) WT and mutant MEFs and those rescued by transiently expressing Wdr35::EmGFP serum starved for 24h hours, fixed and stained with acetylated *α* tubulin (green) and *γ* tubulin (magenta), nuclei (blue). Boxed regions are enlarged below, and arrows point at ciliary axoneme stained for acetylated *α* tubulin. (B) Quantification of cilia length for acetylated *α* tubulin. n= total number of cells from three different biological replicates (represented by different shapes). Asterisk denotes significant p-value from t-test: (*, P<0.05), (**, P<0.01), (***, P<0.001). (C) Percentage of acetylated *α* tubulin positive ciliated cells. (D) 24 hrs serum starved WT and mutant MEFs stained for nuclei (blue), acetylated *α* tubulin/ polyglutamylated tubulin (green), rootletin (cyan) and transition zone proteins MKS1/NPHP-1 (magenta) show no difference in the localization of transition zone protein MKS1 and NPHP-1. Grey scale enlarged regions are labeled green (G), magenta (M), and cyan (C). (E) TEM micrographs of 300 nm sections showing a reduction in overall length of mutant cilia compared to WT MEFs. Scale bar, A, D = 5 *µ*m and E= 1 *µ*m. **Figure 1–Figure supplement 1. *Wdr35***^−/−^ **and *Dync2h1***^−/−^ **mutant cells have a drastic reduction in cilia length but have no difference in the number of cilia**.

### Wdr35 null cells have intact IFT-B complexes with a retrograde defect and unstable IFT-A holocomplexes which fail to enter cilia

Axoneme elongation during cilia assembly requires the import of key building blocks from their place of synthesis in the cell body via IFT. We focused first on the anterograde IFT machinery. We found that IFT-B complex proteins have similar retrograde trafficking defects in both *Wdr35*^−/−^ and *Dync2h1*^−/−^ cells (**Figure 2A, B**), accumulating beyond the length of the acetylated axoneme. We next looked to see if IFT-B complex assembly is disturbed in the absence of WDR35 by endogenous immunoprecipitation (IP) of IFT88 followed by mass spectrometry (MS). IFT88 is the link between peripheral and core IFT-B complexes (**Figure 2C**), interacting with IFT38 on the peripheral side and IFT52 on the core side ***(Katoh et al., 2016; Mourão et al., 2016)***. MS analysis of immunoprecipitates from E11.5 *Wdr35*^+/+^ and *Wdr35*^−/−^ embryo lysates revealed no difference in stoichiometric composition of IFT-B complexes (**Figure 2D, E**), suggesting only the increased ciliary distribution from pools at the ciliary base due to failed exit from the mutant cilia. We were able to isolate the entire IFT-B complex (14 out of 16 IFT-B components) aside from IFT70, which is not yet reported in mouse as well as phosphoprotein IFT25, which binds as a heterodimer to IFT27 ***(Bhogaraju et al., 2011; Funabashi et al., 2017; Katoh et al., 2016; Wang et al., 2009)*** and is necessary for Hh signaling ***(Keady et al., 2012)***. Thus, there is no defect in entry of IFT-B holocomplexes into cilia, but their ability to exit from cilia is impaired in the absence of WDR35.

**Figure 2.**
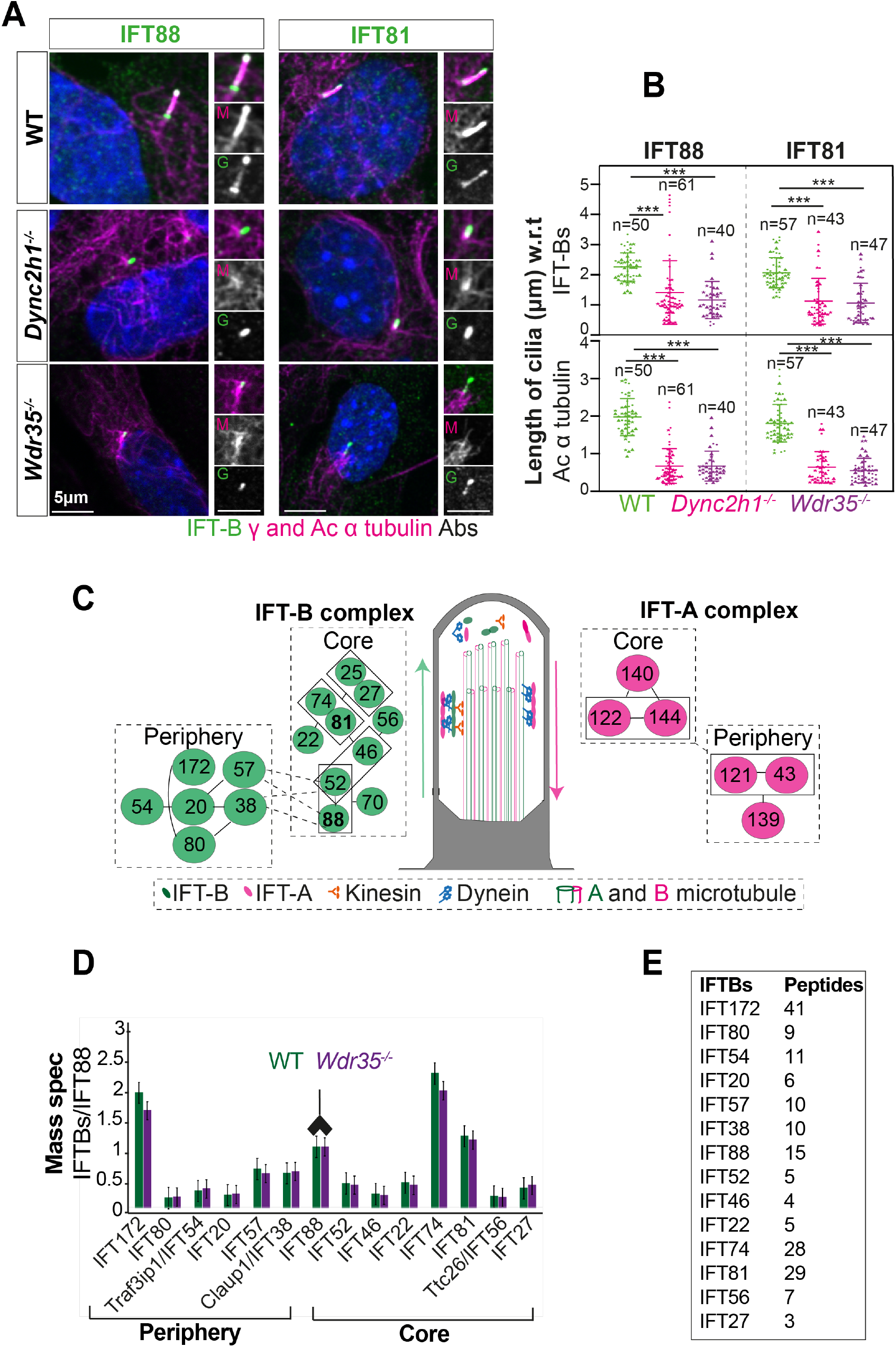
Wdr35^−/−^ cilia exhibit retrograde transport defects of IFT-B, similar to Dync2h1^−/−^, although IFT-B complex assembly is unaffected. (A) IFT-B (green) accumulates beyond the axoneme (Ac-*a* tubulin, magenta) in *Wdr35*^−/−^ and *Dync2h1*^−/−^ cilia from 24 hr serum-starved and fixed MEFs. (B) Length quantification shows IFT-B accumulates beyond acetylated *a* tubulin in significantly shorter mutant cilia. n= total number of cells from three different biological replicates represented by different shapes. Asterisk denotes significant p-value from t-test: (*, P<0.05), (**, P<0.01), (***, P<0.001). Scale bars = 5 *µ*m. (C) Schematic of intraflagellar transport (IFT) pathway in cilia. (D-E) Despite differences in localization, IFT88-IP/MS analysis of E11.5 WT and *Wdr35*^−/−^ littermate embryos reveal no difference in the composition of the IFT-B complex. Antibody highlights bait (IFT88) for IP. (D) Normalized LFQs to IFT88 intensity reveals no difference between WT and *Wdr35*^−/−^ IFT-B complex composition. N= 4 embryos/genotype. (E) The number of unique peptides identified in IP-MS.

We next checked the stability of the IFT-A holocomplex by endogenous immunoprecipitation of IFT-A core protein IFT-140 and its interactors. Whilst IFT140 immunoprecipitated all six components of the IFT-A complex from *Wdr35*^+/+^ embryo lysates, in *Wdr35*^−/−^ samples both peripheral components IFT139 and IFT43 were missing in MS (**Figure 3A**) and immunoblot (**Figure 3B**). Moreover, core components were also significantly reduced in the purified mutant complex, suggesting that WDR35 is critical for the stable IFT-A complex assembly. To distinguish defects in stability from defects in assembly, we looked at total IFT-A component protein levels and we found total IFT139 and IFT43 levels were also undetectable on blots with lysates from both *Wdr35*^−/−^ MEFs (**Figure 3C, E**) and embryos (**Figure 3D, F**). This suggests WDR35 is not only critical for the formation of stable IFT-A holocomplex but is also required for stability of its peripheral components. In contrast, the core components of the IFT-A complex were nearly equally expressed in both WT and *Wdr35*^−/−^ lysates except for IFT122, which had higher expression levels in *Wdr35*^−/−^ compared to control (**Figure 3D, F**). Other core components have been shown to have higher levels in the absence of WDR35 in human fibroblasts ***(Duran et al., 2017)***. Our work supports previous studies demonstrating interdependent protein expression of IFT-As, which might be required for their coordinated function ***(Behal and Cole, 2013; Duran et al., 2017; Fu et al., 2016; Picariello et al., 2019; Zhu et al., 2017)***.

**Figure 3.**
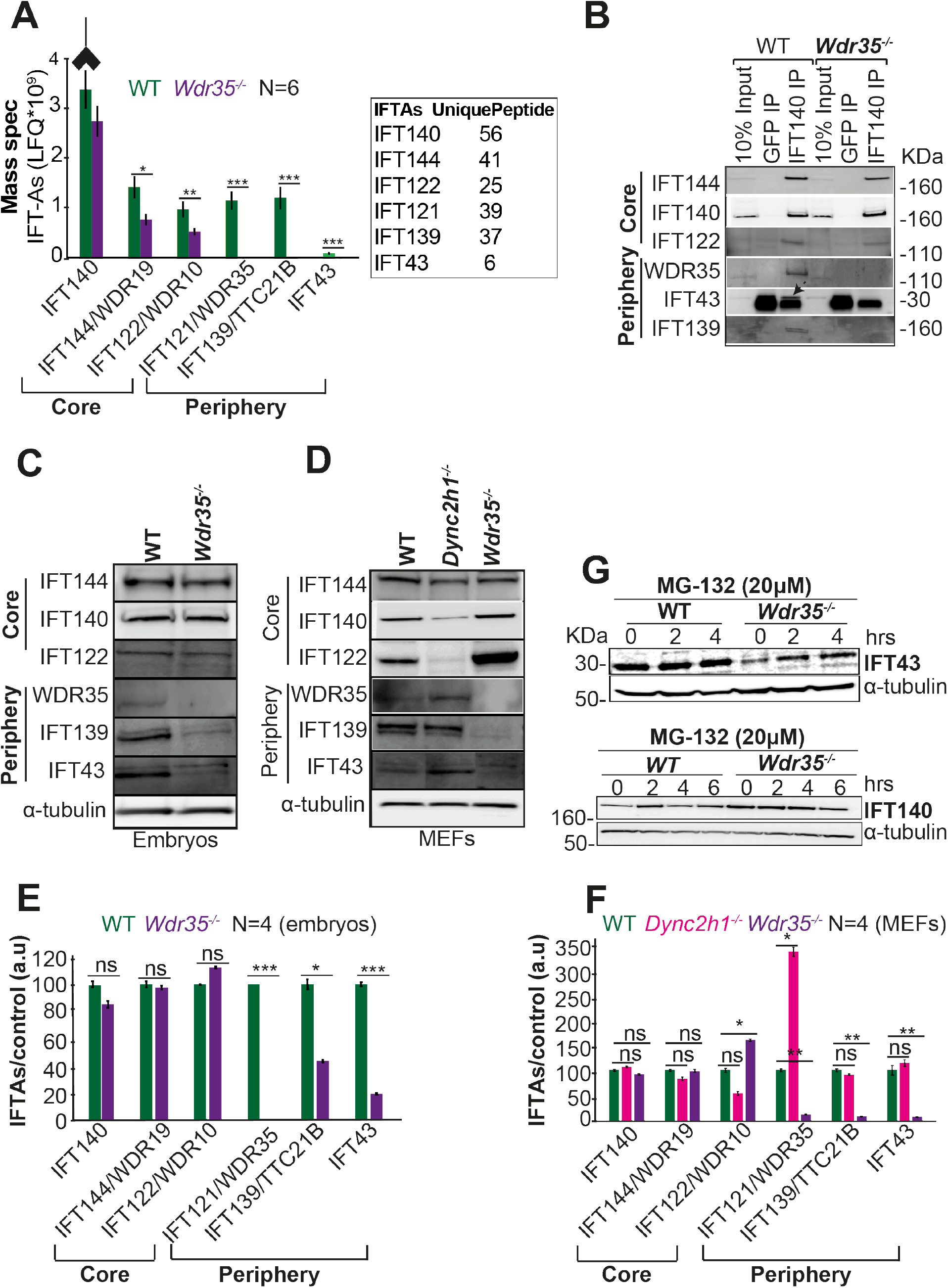
WDR35 is essential for the stability of the IFT-A complex. (A) IP/MS data shows the stability of the IFT-A complex is disrupted in *Wdr35*^−/−^lysates. N=6 embryos per genotype. Antibody highlights bait (IFT140) for IP. (B) Immunoblots confirm the peripheral IFT-A complex is unstable in *Wdr35*^−/−^ mutants. IFT43 runs close to the molecular weight of IgG, is shown by an arrow as IFT43 band over the IgG band from IFT140 IP in WT. The corresponding band is absent in *Wdr35*^−/−^ lysates. (C-D) Immunoblots for the total level of IFT-A subunits in (C) E11.5 embryo and (D) MEF lysates show peripheral components IFT139 and IFT43 to be missing in Wdr35 mutant cells, quantified by densitometry in (E) embryos and (F) MEFs. N= biological replicates. Asterisk denotes significant p-value from t-test: * P< 0.05, ** P<0.01, *** P<0.001. (G) Inhibition of the proteasome by treatment with MG-132 rescues IFT43 stability in *Wdr35*^−/−^ MEFs.

These results suggest WDR35 might be a link between IFT-A core and periphery proteins, required to protect them from degradation. To investigate this further, control and mutant MEFs were treated with 20µM MG-132 proteasome inhibitor (**Figure 3G**). Treated cells rescued expression of IFT43, which suggests that in the absence of WDR35, some peripheral proteins may be targeted by the proteasomal degradation pathway. Interestingly IFT139 and IFT121 are degraded in IFT43 null cells and both are rescued similarly by MG-132 treatment ***(Zhu et al., 2017)***, confirming that the stability of IFT-A complex proteins is interdependent.

We next looked at the localization and levels of the IFT-A components by immunofluorescence. All IFT-A components accumulated in *Dync2h1*^−/−^ cilia suggesting entry of IFT-A holocomplexes is not affected, but return from the distal tip cannot occur in the absence of the dynein motor (**Figure 4A, B**). In contrast, in *Wdr35*^−/−^ MEFs, IFT-A core components fail to enter cilia and remain restricted to the transition zone (**Figure 4A**), as shown by the difference in length covered by IFT-A components relative to cilia length measured by acetylated tubulin staining (**Figure 4B**), whereas peripheral proteins were undetectable (**Figure 4A, B**), consistent with degradation (**Figure 3C-F**). These results are consistent with previous reports of the inter-dependence of IFT-A components for ciliary localization. IFT140 is decreased in cilia of *IFT122* mutants in mouse and fly ***(Lee et al., 2008; Qin et al., 2011)***, IFT139 is reduced in the flagella of *Chlamydomonas* with *IFT144* mutation ***(Iomini et al., 2009)***, and IFT144 fails to get recruited into cilia in *WDR35*^−/−^ RPE cells ***(Fu et al., 2016)***. IFT-A proteins require CPLANE chaperones for holocomplex assembly and cilia entry ***(Toriyama et al., 2016)***. In all cases, failure of IFT-A holocomplex integrity impairs its recruitment into the cilia axoneme. Recent cryo-EM work had suggested IFT-A is being carried by IFT-B trains inside the *Chlamydomonas* flagella in WT cells and these structures are missing in the *IFT139* mutant ***(Jordan et al., 2018)***. Our work in the mammalian system in the absence of WDR35 has a similar effect with IFT-B proteins accumulating inside the cilium whilst IFT-A core proteins accumulate proximal to the cilia base, and the peripheral components are degraded.

**Figure 4.**
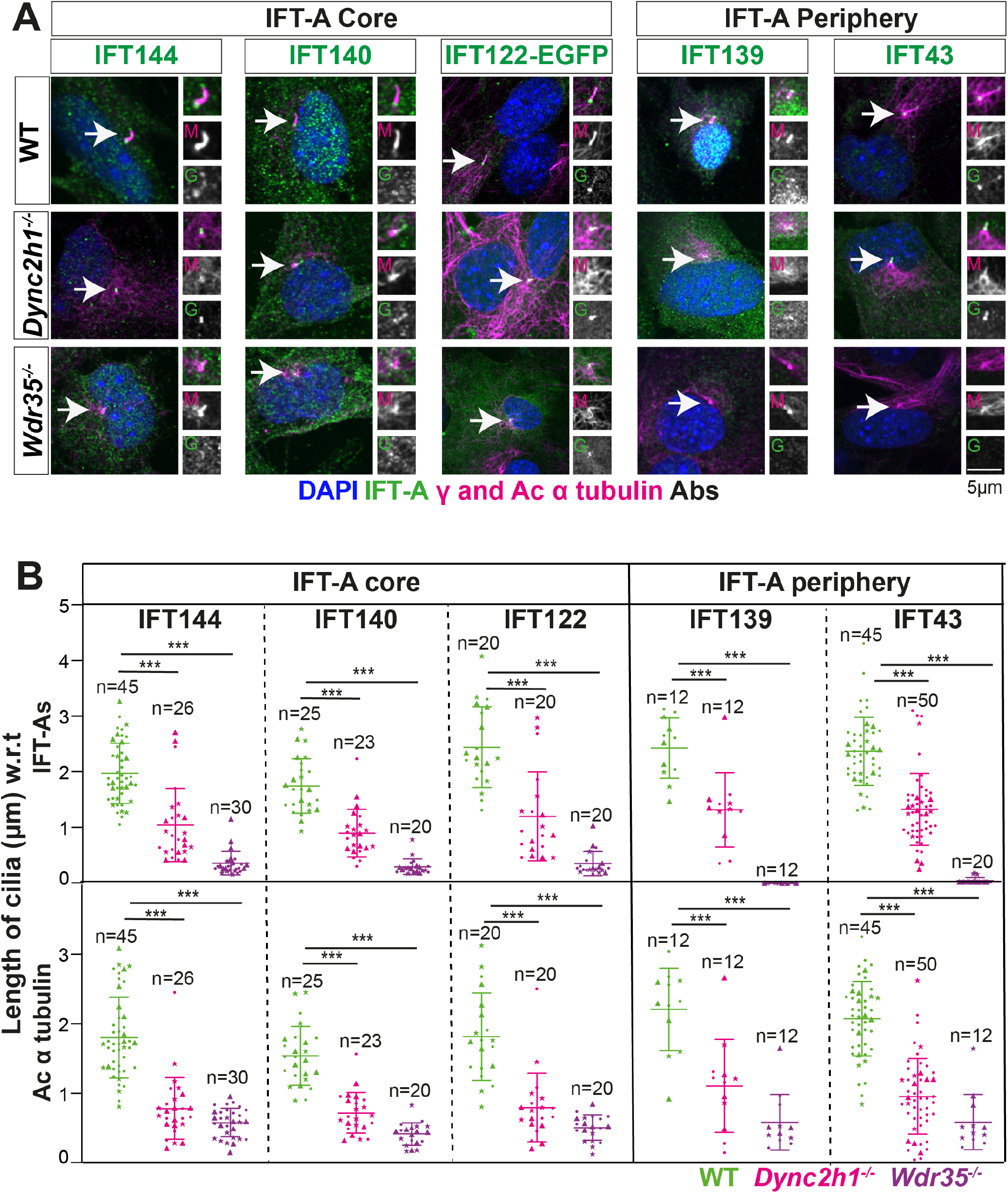
WDR35 is critical for the recruitment of IFT-A components into cilia. (A) MEFs serum starved for 24h reveal a retrograde transport defect in *Dync2h1*^−/−^ versus a failed recruitment of IFT-A proteins into *Wdr35*^−/−^. Cells are fixed and stained for respective IFT-A (green) and *r* and acetylated *a* tubulin (magenta). Due to a lack of specific immunoreagents, IFT122 signal is from transiently expressed Ift122::GFP. All other panels represent endogenous signal detected by IF. Arrows point at cilia. Scale bars = 5 *µ*m. (B) Quantification of cilia length for acetylated *a* tubulin, and IFT-As. n= total number of cells from three different biological replicates (represented by different shapes). Asterisk denotes significant p-value from t-test: (*, P<0.05), (**, P<0.01), (***, P<0.001).

### Membrane proteins fail to be recruited into Wdr35^−/−^ cilia

Cilia membrane protein cargos must be synthesized in the cell body and trafficked into cilia through a variety of direct and indirect routes. These include lateral diffusion ***(Leaf and Von Zastrow, 2015; Milenkovic et al., 2009)***, recycling of membrane proteins via the endocytic pathway (Boehlke et al., 2010) as well as directly from Golgi-derived vesicles ***(Follit et al., 2008, 2006; Kim et al., 2014; Witzgall, 2018)***. Moreover, cilia membrane content is dynamically regulated in response to signals. We investigated endogenous levels and localizations of small membrane associated GTPases ARL13B and ARL3, which are constitutively localised in control cilia (**Figure 5A**) We saw that while they accumulate in excess in *Dync2h1*^−/−^ mutants as per a retrograde defect, strikingly they fail to be recruited to *Wdr35*^−/−^ cilia. Similarly, we also tested appropriate dynamic recruitment of the GPCR co-receptor Smoothened (SMO), which is recruited to the cilia in response to Hh ligand. SMO is not excluded from *Dync2h1*^−/−^ mutant cilia in the absence of Hh. In contrast, even in the presence of Hh activation, SMO fails to enter *Wdr35*^−/−^ cilia. Detecting low levels of endogenous protein localization and their mislocalization in *Wdr35* mutants by immunofluorescence can be challenging. To overcome this, we transiently transfected membrane cargos, including fluorescently-tagged ARL13B and SMO (**Figure 5B and Movie 2**), which are effectively trafficked to the cilia of control cells. However, they fail to localize to *Wdr35*^−/−^ cilia, with some accumulation at the cilia base. Interestingly, overexpressed ARL13B, when not transported to cilia, is concentrated on other membranes, particularly the plasma membrane, whereas SMO was predominantly localized in vesicles in the cytoplasm of mutant cells (**Figure 5B and Movie 2**). Similarly, few other ciliary membrane proteins like EVC1/2, INPP5E, SSTR3, 5HT6, and MCHR-1 have been shown to have failed ciliary localisation in *WDR35* mutants ***(Caparrós-Martín et al., 2015; Fu et al., 2016)***.

**Figure 5.**
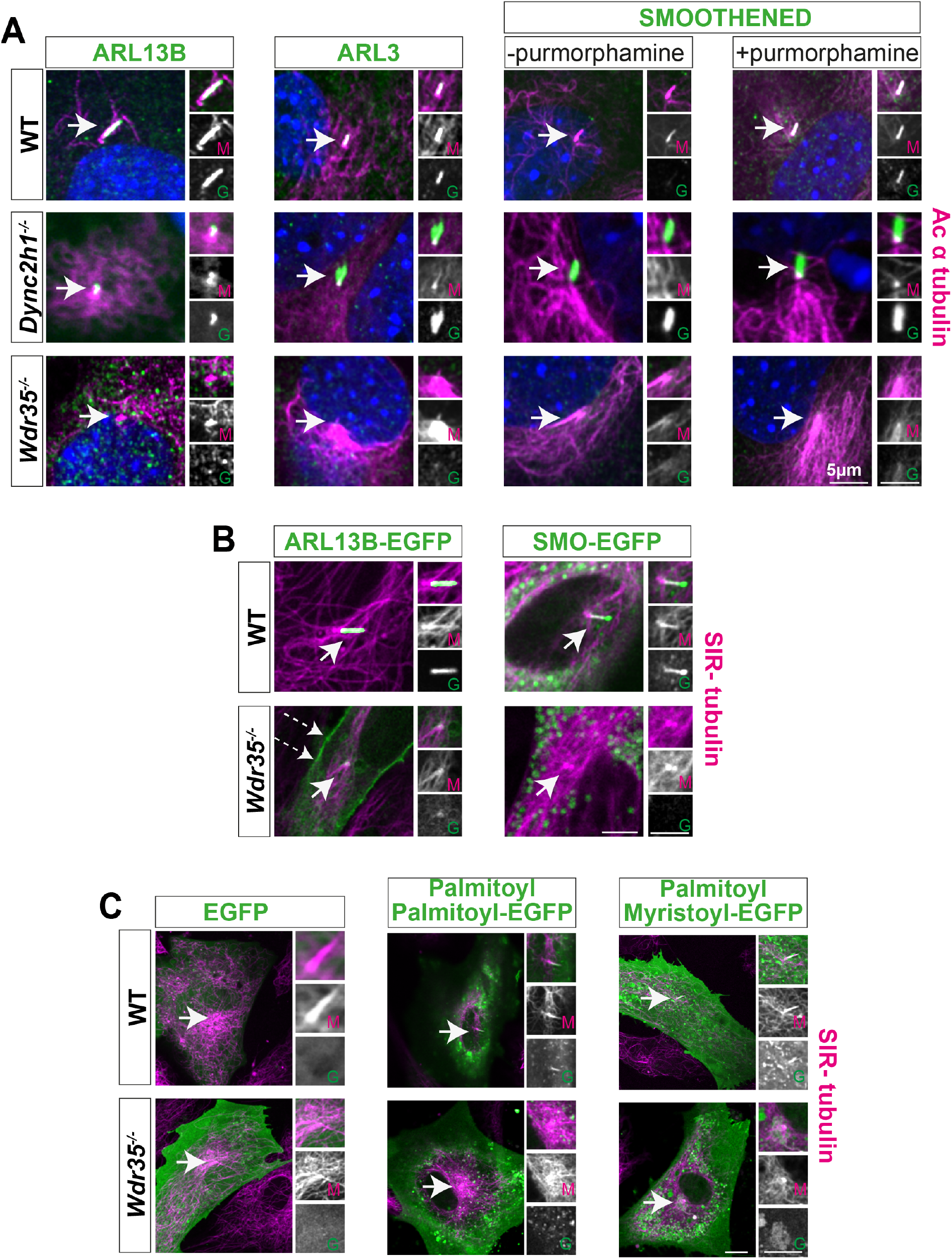
Membrane proteins fail to localise to Wdr35^−/−^ cilia. A) 24 hr serum-starved WT, *Wdr35*^−/−^ and *Dync2h1*^−/−^ MEFs stained for ARL13B, ARL3, and Smoothened (green), and acetylated *a* tubulin (magenta) show failed localization of membrane proteins in *Wdr35*^−/−^ and retrograde transport defect in *Dync2h1*^−/−^. (B) ARL13B-EGFP and Smoothened-EGFP (green) expressing ciliated cells stained with SiR-tubulin (magenta) show failed localization of exogenously expressed membrane proteins inside mutant cilia **(Movie 2)**. Dashed arrows point at the enrichment of ARL13B on the membrane in the mutant. (C) 24 hrs serum-starved cells expressing respective general lipidated GFP cargos (green) and stained for SiR-tubulin show enrichment of lipidated GFP in WT cilia and failed localization in the mutant. Arrows point at cilia in all images. Scale bars 5 *µ*m.

In trypanosomes, localization of flagellar membrane proteins was shown to be dependent on lipid rafts highly enriched in axonemes ***(Tyler et al., 2009)***. Here, dual acylation was shown to direct potential association with lipid rafts, membrane microdomains that function as specialized platforms for protein/lipid transport and signaling. Similarly, in mammalian cells, palmitoylation of ARL13B was required for cilia entry and maintenance ***(Roy et al., 2017)***. We asked about the ability to recruit general lipidated cargo in *Wdr35*^−/−^MEFs. We examined the localization of lipidated motifs tagged to EGFP ***(Williams et al., 2014)*** to look at specialized membrane microdomains. In WT MEFs, untagged EGFP is present in the cell, but not in the cilium. When tagged with either myristoylation and palmitoylation (MyrPalm) or dual palmitoylation (PalmPalm) motifs, EGFP robustly enriches within cilia (**Figure 5C**). We observed no enrichment of dual geranylation (GerGer) modified EGFP within control fibroblast primary cilia (data not shown), in contrast to the low level expression previously reported in the most proximal portions of highly specialized olfactory sensory cilia ***(Williams et al., 2014)***. This suggests that cell-type and cilia-specific differences exist. In contrast, in *Wdr35*^−/−^ MEFs, both the myristoylation and palmitoylation (MyrPalm) or dual palmitoylation (PalmPalm) eGFP accumulated around the basal body (**Figure 5C**). This failure to recruit lipidated cargoes into *Wdr35* mutant cilia is consistent with a more global trafficking disruption of ciliary-destined membrane-microdomains, containing broad categories of the membrane and membrane-associated cargos.

### WDR35 and other IFT-A complex proteins share close sequence and structural similarity to COPI complex proteins α and β′

It has previously been suggested that IFTs have evolved from a protocoatomer ***(Jékely and Arendt, 2006; Taschner et al., 2012; van Dam et al., 2013)***. Three classic coat complexes (COPI, COPII and clathrin) exist which perform similar functions but on different membranes and follow different routes through the cell. They are made of different protein components, which share a similar division of labor characterized functionally as either adaptors or cage forming proteins. Although components like the cage proteins share significant structural homology in organization of protein domains, they do not share detectable sequence homology ***(Field et al., 2011)***. In the case of IFT172, the N-terminal WD repeat domain was necessary and sufficient for membrane binding in vitro ***(Wang et al., 2018)***. Given the defects in ciliary membrane content observed in the *Wdr35* mutant cilia, we hypothesized that WDR35, in collaboration with other IFT-A complex proteins may be required for moving ciliary membrane cargos between donor membranes, such as the Golgi or the ciliary pocket, to their destination ciliary membrane. WD40 repeat (WDR) and tetratricopeptide repeat (TPR) motifs are common throughout cellular proteomes and are involved in a wide range of biological processes. Agnostic of structure, we used deep sequence analysis and homology modeling of the whole human proteome to ask what proteins were most similar to IFT-A components. Simple alignment strategies with proteins such as these IFTs, which contain tandem repeat motifs, could erroneously align with other repeat proteins to suggest a close evolutionary relationship where none exists (i.e., false positives). To address this, we used four IFT-A subunits (IFT144, IFT140, IFT122 and IFT121) and two of IFT-B (IFT80 and IFT172) as seed sequences for multiple iterative rounds of homology searches via profile-HMM alignment ***(Remmert et al., 2011)***. We then clustered the resulting proteins based on sequence similarity, as previously described ***(Wells et al., 2017; Wells and Marsh, 2019)***. This was repeated using the COP proteins as seeds for reverse analysis. Together, these reciprocal analyses revealed that out of the entire proteome, COPI *a* and *β*′ cluster most closely with 6 IFT proteins (two IFT-B and four IFT-A components), both having TPR and WD40 repeats (**Figure 6A**). In contrast, homology searches with COPI */* and COPI *γ*1/2, which have HEAT/ARM repeats, did not yield any IFT components, as was the case with COPI *ε*, which has TPR repeats but no WD40 domains. COPI *δ* and COPI *γ*1/2, which have no identifiable repeat domains, are most closely related to adaptor protein complex subunits AP2 and AP3. In summary, using multiple rounds of sequence homology searches, we generated a broad set of putatively related repeat proteins, clustering of which reveals clear relationships between coatomers and IFT-A/B complex components.

**Figure 6.**
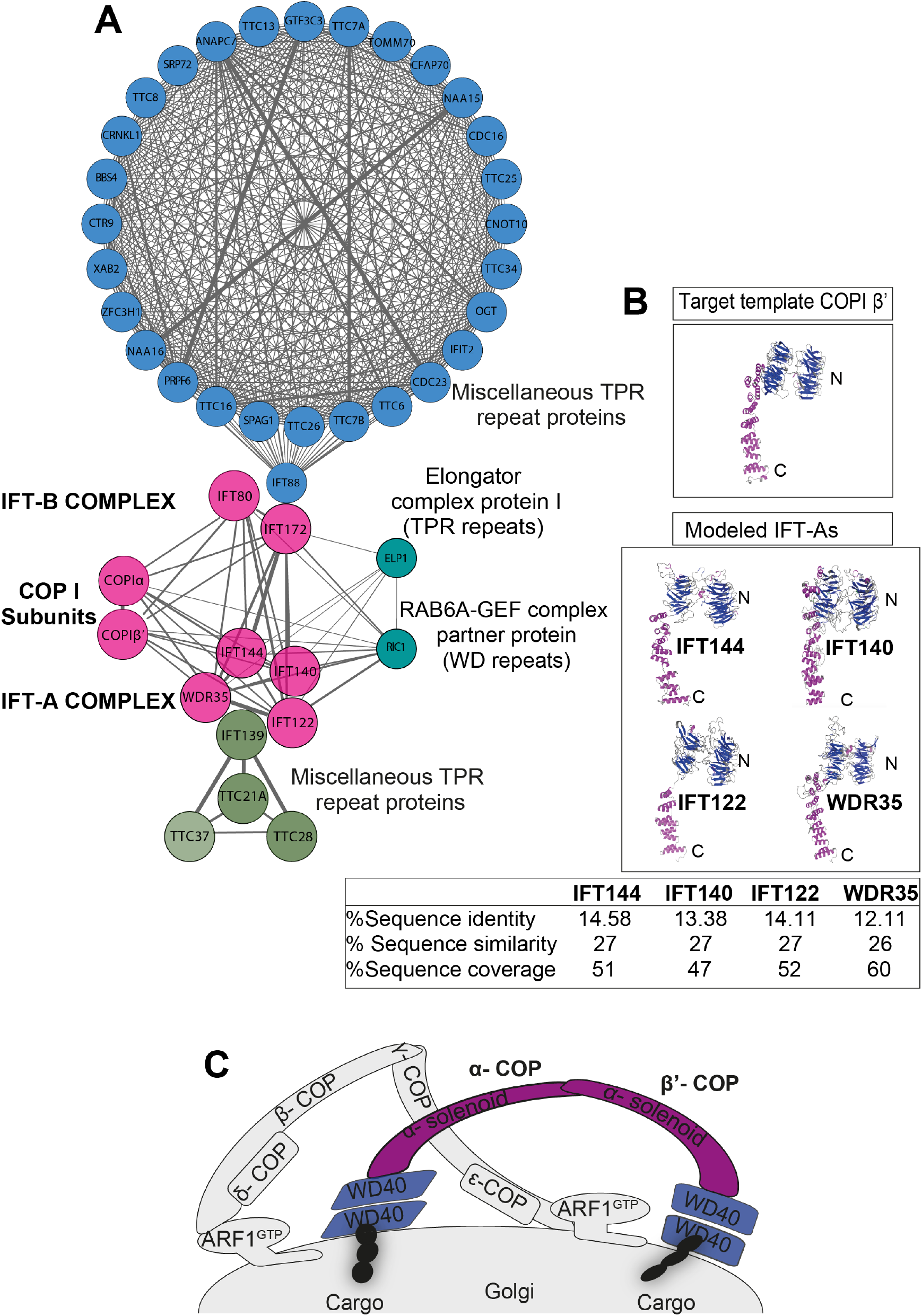
IFT-A subunits have close sequence and structural similarity to (α and /’) COPI subunits. (A) Clusters of IFT and COPI subunits generated from the results of reciprocal sequence similarity searches with HHBlits using IFT144, IF140, IF122, and WDR35 as initial search queries, suggest a very close similarity between a subset of IFT proteins and the COPI (*a* and *β*′) subunits. Clusters are color-coded according to protein structural motifs with TPR repeat proteins (blue) and dual WD40 repeat and TPR repeat-containing proteins (magenta). Lines between clusters indicate sequence-based proximity. (B) Structure prediction showed IFT144, IFT140, IFT122, and WDR35 to have close structural similarity to COPI complex proteins (*a* and *β*′). 2.5Å X-ray structure of *β*′ (PDB:3mkq) and IFT-A proteins are shown with N-terminal WD40 repeat (blue) and C-terminal TPR repeats (magenta). Sequence identity, similarity, and coverage between COPI -*β*′ and respective IFT-A proteins are shown in the table below. (C) The cartoon representation of COPI coat assembly subunits *α, β, β*′, *γ, ϵ*, and *δ*. The *α* and *β*′ subunits have WD40 repeat (blue) and TPR repeat (magenta), and WD40 hydrophobic domain insertion in the Golgi membrane facilitates membrane curvature initiating COPI coat assembly.

Next, we used SWISS-MODEL ***(Waterhouse et al., 2018)*** to predict the structures of IFT-A proteins. COPI *α* and *β*′ structures were top hits with 12%-15% sequence identity and 26%-27% sequence similarity to four IFT-A complex proteins (IFT144, IFT140, IFT122, and WDR35). Based on the target-template alignment models are built using ProMod3, ribbon diagrams of all these 4 IFT-A subunits modeled structures with two N-terminal seven-bladed WD40 *β* propellers and C terminal extended TPR repeats, also found in *α* and *β*′ COPI proteins (**Figure 6B**), as previously modeled for WDR35 ***(Mill et al., 2011)***. The remaining 2 IFT-A subunits were not possible to model accurately. IFT139 contains only TPR repeats with limited sequence similarity to the *ε* subunit of COPI coatomer ***(van Dam et al., 2013)***. IFT43 is the smallest and unstructured protein and could not be modeled and is presumed to be made of *α*-helices ***(Taschner et al., 2012)***. While undertaking this work, a crystal structure for IFT80 was published highlighting that while it had the same domain organization, IFT80 adopted an altered 3D configuration of the second - propeller domain from *β*′-COP and also formed a dimer unlike the triskelion COP I cage ***(Taschner et al., 2018)***. However, while not a solved structure, purified IFT172 adopted two configurations by negative stain electron microscopy (EM) when incubated with and without lipids, the former being mutually restrictive with IFT57 binding ***(Wang et al., 2018)***. However, respecting the limitations of homology modeling without solved structures, we found 4 IFT-A proteins (IFT144, IFT140, IFT122, and IFT121) have very high sequence and structural similarity to COPI *α* and *β*′ subunits with N-terminal WD40 repeats and C-terminus TPR region (**Figure 6B**).

### Distinct ultrastructural ciliary defects are observed between disruption of IFT-A versus the retrograde IFT motor

Short cilia are difficult to resolve by light microscopy. We undertook ultrastructural studies to examine phenotypes with higher resolution in MEFs. In all genotypes, ciliation was observed to start very close to the nucleus and remains close to the Golgi stacks throughout cilia elongation (**Figure 7A, Movie 3; Figure 7B, Movie 4; Figure 7C, Movie 5; Figure 7 Supplement 2, Movie 6**). In control MEFs, even after 24 hours of serum starvation, only 1% of cilia were observed to emerge from the cell, highlighting the deep-seated ciliary pocket in MEFs (**Figure 7B, Movie 4; Figure 7 Supplement 1 and 3A**), as described for RPE-1 cells ***(Molla-Herman et al., 2010)***. In control MEFs, polymerised microtubules formed a well-structured axoneme (**Figure 7B, Movie 4; Figure 7 Supplement 1 and 3A**) as previously described in MEFs ***(Rogowski et al., 2013)*** and reported in other primary cilia ***(Kiesel et al., 2020; Molla-Herman et al., 2010)***. Additionally, microtubules can be seen attached at the cilia base and radiating in different directions in the cell (**Figure 7 Supplement 1**). In contrast to the well-defined ciliary membrane and well polymerized microtubules of the control axoneme, *Wdr35*^−/−^ cilia have ‘wavy’ membranes and disorganized microtubules (**Figure 7C, movie 5 and Figure 7 Supplement 3B**). Mammalian *Dync2h1*^−/−^ mutants retained a well-defined ciliary membrane and an apparently well-structured axoneme present throughout (**Figure 7 Supplement 3C**), similar to previous reports of the *la14* dynein mutant in *Chlamydomonas* ***(Pigino et al., 2009)***. Stacked standing trains with a periodicity of 40nm were reported in *la-14* mutants ***(Pigino et al., 2009; Stepanek and Pigino, 2016)*** and in our *Dync2h1*^−/−^ mutant axonemes, we observed similar stacking of stalled IFT trains with a periodicity of 40 nm, irrespective of the length of mutant cilia (**Figure 7 Supplement 3C** and ***(Liem et al., 2012)***). Although IFT-Bs also accumulated in *Wdr35*^−/−^ cilia (**Figure 2A, B**), these stripes were not observed (**Figure 7C, Movie 5; Figure 7 Supplement 3B**) suggesting that both IFT-B and IFT-A are required to form the higher ordered IFT trains which stall in *Dync2h1*^−/−^ mutants.

**Figure 7.**
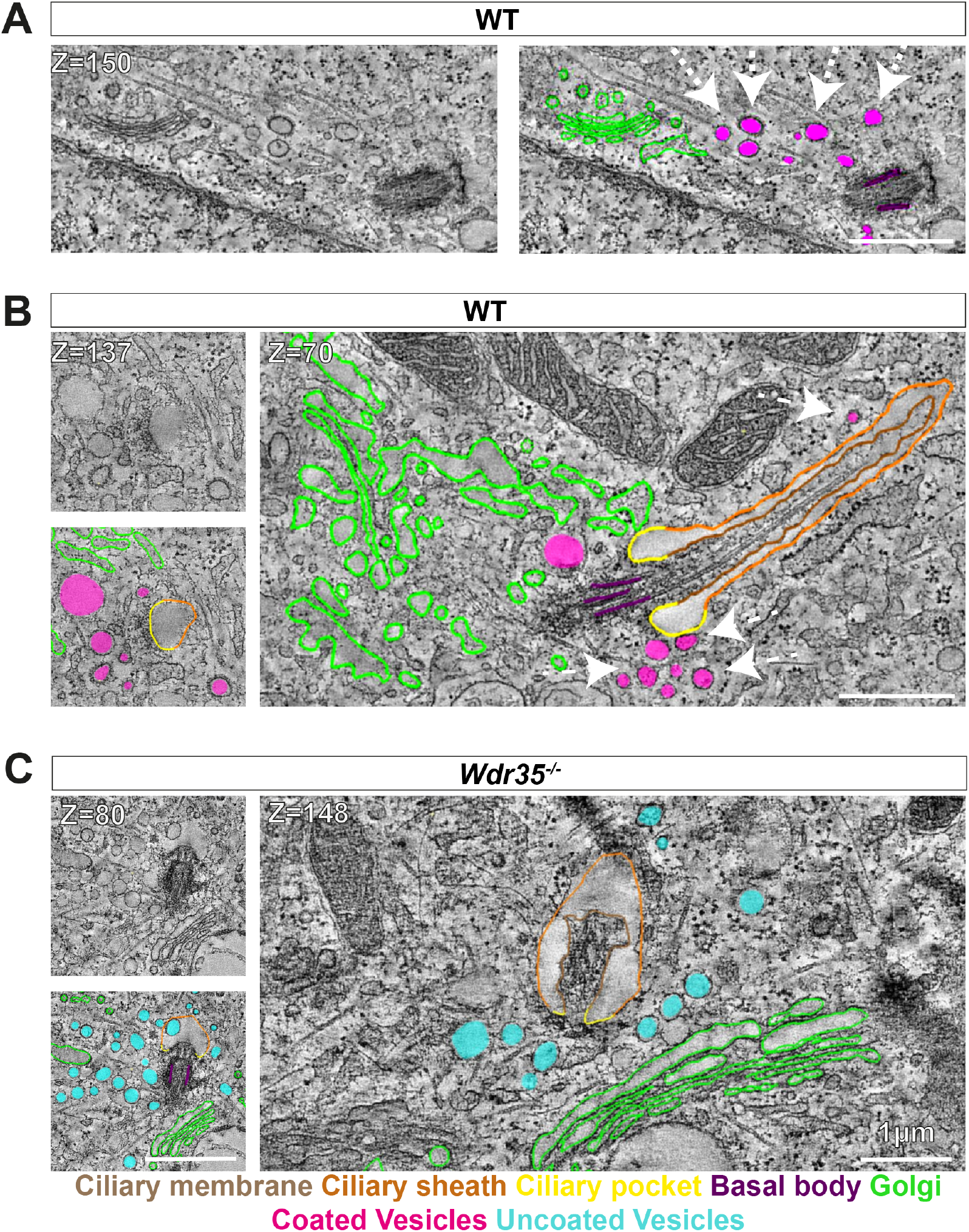
Electron-dense vesicles are observed tracking between the Golgi and cilia base in WT fibroblasts whereas a ‘coat-less’ vesicles accumulate around Wdr35 mutant cilia. The tilt series of TEM samples were made from 24 hr serum starved MEFs. Reconstructed tomograms are color-coded to highlight the ciliary membrane (brown), ciliary sheath (orange), ciliary pocket (yellow), basal body (purple), Golgi (green), electron-dense coated vesicles (magenta), and vesicles lacking electron cloud (cyan). (A) Z-stacks from 600 nm TEM serial tomograms of WT MEFs show track of electron-dense vesicles between the Golgi and cilia **(Movie 3)**. Dashed arrows point at the path of vesicles between the Golgi and cilia. The image in the left panel is segmented in the right panel. (B) Z-stacks from 300 nm tomograms from WT MEFs show electron-dense coated vesicles close to the cilia base and along the length **(Movie 4)**. Dashed arrows point at coated vesicles near cilia. (C) Z-stack from 600 nm serial tomogram from *Wdr35*^−/−^ MEFs has a massive accumulation of vesicles in a 2µm radius of the cilia base (cyan), and these vesicles lack a visible coat, or electron-dense cloud on them **(Movie 5)**. The length of cilia is drastically reduced, the ciliary membrane is wavy, and axoneme microtubules are broken in the mutant. (B and C) On left is the same Z-stack in the upper panel segmented in the lower panel, and on the right is another Z-stack from the same tomogram. Scale bars = 1*µ*m.

### WDR35 forms vesicular coats to trafic membrane proteins packed in vesicles from the Golgi to cilia

We further tested our hypothesis for the coatomer function of IFT-As by transmission electron microscopy (TEM) analysis of ciliated MEFs. We observed electron-dense coated vesicles between the Golgi and cilia in WT MEFs (**Figure 7A, Movie 3**). All TEM results showed the Golgi to be in close vicinity of the cilium (**Figure 7A, Movie 3; Figure 7B, Movie 4; Figure 7C, Movie 5; Figure 7 Supplement 2, Movie 6**). We also observed these coated vesicles clustering at the cilia base (**Figure 7B, Movie 4**) and bulging from ciliary pockets and ciliary sheaths in WT MEFs (**Figure 7 Supplement 1**). These electron-dense vesicles around control cilia were more prominent at the early stage of ciliogenesis in EM (**Figure 7A, Movie 3**).

In contrast, in *Wdr35*^−/−^ mutant cells, there is a ten-fold increase in the average number of vesicles around the cilia base (**Figure 7C, Movie 5; Figure 7 Supplement 2, Movie 6; Figure 7 Supplement 3B; Figure 8, Figure 8 Supplement 1A**). Importantly, virtually all of these mutant vesicles lack the electron-dense coats observed in control cells (**Figure 7C, Movie 5; Figure 7 Supplement 2, Movie 6; Figure 7 Supplement 3B; Figure 8, Figure 8 Supplement 1A**). Notably, we did observe other electron-dense coats, likely clathrin, on budding vesicles at the plasma membrane in these same *Wdr35*^−/−^ cells, emphasizing that other coats are preserved in these conditions (**Figure 7 Supplement 2, Movie 6, Figure 8B**). Moreover, no difference in the density or distribution of periciliary clathrin-positive vesicles is observed around the base of *Wdr35*^−/−^ mutant cilia (**Figure 8 Supplement 1C, D**). In contrast, the accumulation of uncoated vesicles spreads in a volume 2 µm^3^ around the *Wdr35*^−/−^ ciliary base. In spite of their proximity to the ciliary sheath and their abundance, fusion events were not observed in *Wdr35*^−/−^ mutants (**Figure 7C, Movie 5, Figure 7 supplement 2, and Movie 6**). There is also an increased number of Golgi stacks clustering around the rudimentary cilia in *Wdr35*^−/−^ mutants.

**Figure 8.**
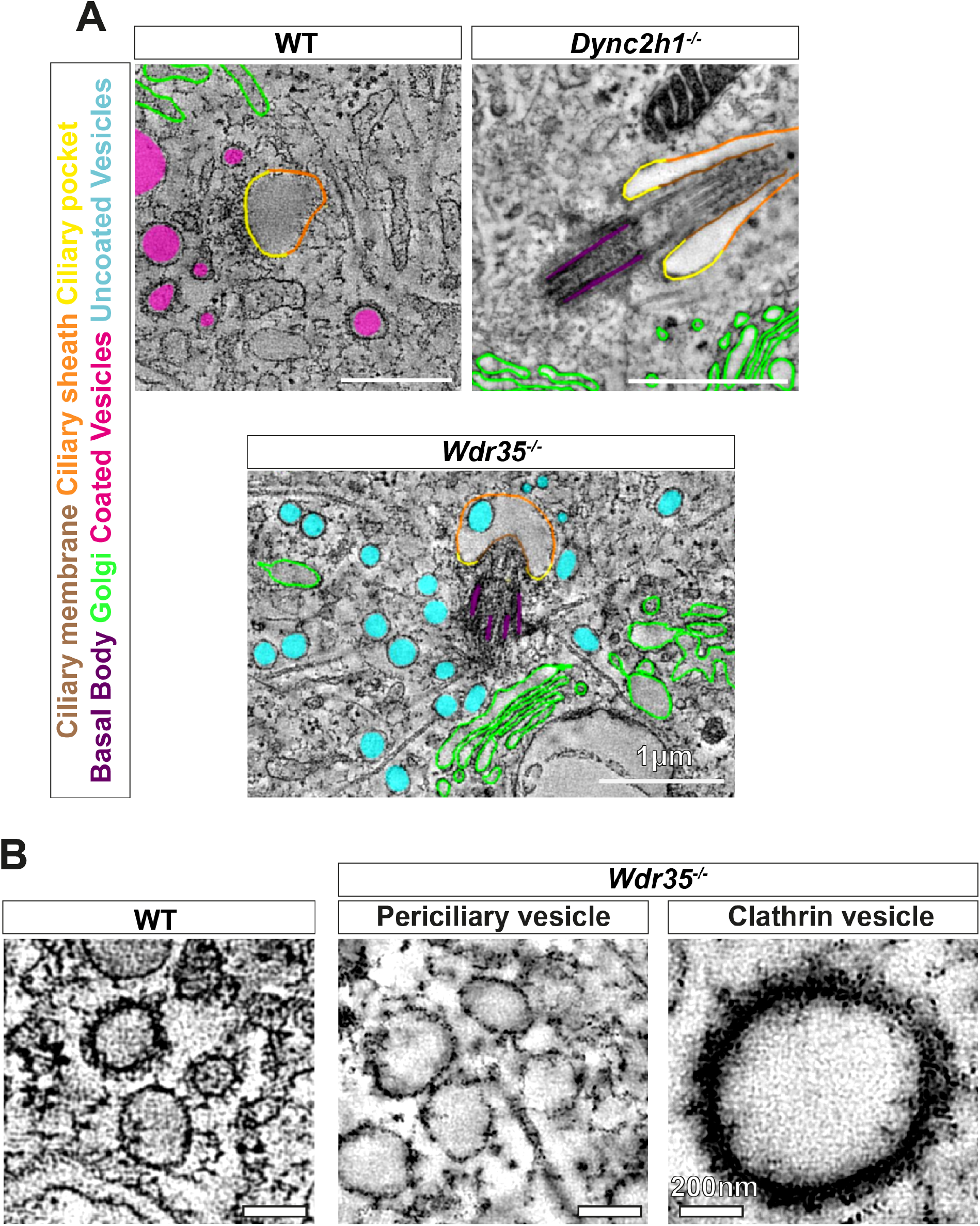
Vesicles clustering around Wdr35 ^−/−^ cilia lack electron dense decorations although electron-dense clathrin coated vesicles are still observed budding from mutant plasma membrane. A) Zoomed-in views of periciliary vesicles observed in WT (**zoomed-Figure 7B, Movie 4**), *Wdr35*^−/−^ (**zoomed-Figure 7C, Movie 5**), *Dync2h1*^−/−^ MEFs 24 hours post-serum starvation show vesicles around WT cilia are coated (magenta) and around *Wdr35*^−/−^ are uncoated (blue). Too few/no vesicles were observed surrounding *Dync2h1*^−/−^ mutant cilia. (B) Enlargement of electron dense coats present around periciliary vesicles in WT **(Movie 4)** and missing in *Wdr35*^−/−^ **(Movie 6)** MEFs. Clathrin vesicles from the same mutant **(Movie 6)** maintain their coat confirming missing electron density on *Wdr35*^−/−^ is not a fixation artefact. Scale bars, A=1 *µ*m and B = 200 nm.

Clathrin-mediated endocytosis at the ciliary base has been previously documented to be regulating export from cilia ***(Molla-Herman et al., 2010)***. To test whether these vesicles might be important for the import or export of cargo from cilia, we analyzed *Dync2h1*^−/−^ cilia, which we showed to be saturated with IFTs (**Figure 2 and Figure 4**) and membrane protein cargo (**Figure 5**) in the absence of retrograde transport. Consistent with the redistribution of IFT pools from the base entirely into the ciliary compartment (**Figure 2 and Figure 4**), we observed no vesicles at the base of *Dync2h1*^−/−^ cilia (**Figure 8A and Movie 7**). Interestingly, ectosomes budding from the tip of *Dync2h1*^−/−^ cilia was more prevalent (**Figure 7 Supplement 3C, Movie 7**), which are previously reported to regulate the content of cilia in a variety of systems ***(Cao et al., 2015; Nager et al., 2017; Wood and Rosenbaum, 2014)***. This strongly suggests that the coated vesicles around the control cilia function to transport cargo from the Golgi to the cilia. In the absence of WDR35, non-coated vesicles accumulate around the ciliary base marking a failure in this process in either the formation and/or maintenance of this coat and subsequent fusion at the target ciliary membrane.

To further confirm our hypothesis that these Golgi-to-cilia vesicular coats are made of WDR35, we performed correlative light and electron microscopy (CLEM) imaging in *Wdr35*^−/−^ MEFs expressing WDR35::EmGFP, which we had previously shown to completely rescue cilia phenotypes (**Figure 1A, B, Figure 9A**). Expressing WDR35::EmGFP in *Wdr35*^−/−^ ensures that every WDR35 particle was labelled with EmGFP, minimizing competition with non-labeled species. Using Airyscan confocal imaging of WDR35::EmGFP MEFs grown on grids for subsequent TEM, we saw WDR35 signal enriched at the ciliary base of rescued mutant cilia. Moreover, we observed that this signal coincided with the re-appearance of electron-dense vesicles now seen by TEM at the cilia base (**Figure 9**). This is reminiscent of the localization reported for IFT140 in photoreceptor ciliary vesicles by EM Gold staining ***(Sedmak and Wolfrum, 2010)***. Together, these results demonstrate that WDR35 is required for the stability of these coated vesicles and that these coated vesicles coincided with WDR35-EmGFP signal, confirming WDR35 supports the assembly of a novel coat on vesicles destined to deliver membrane cargos to cilia.

**Figure 9.**
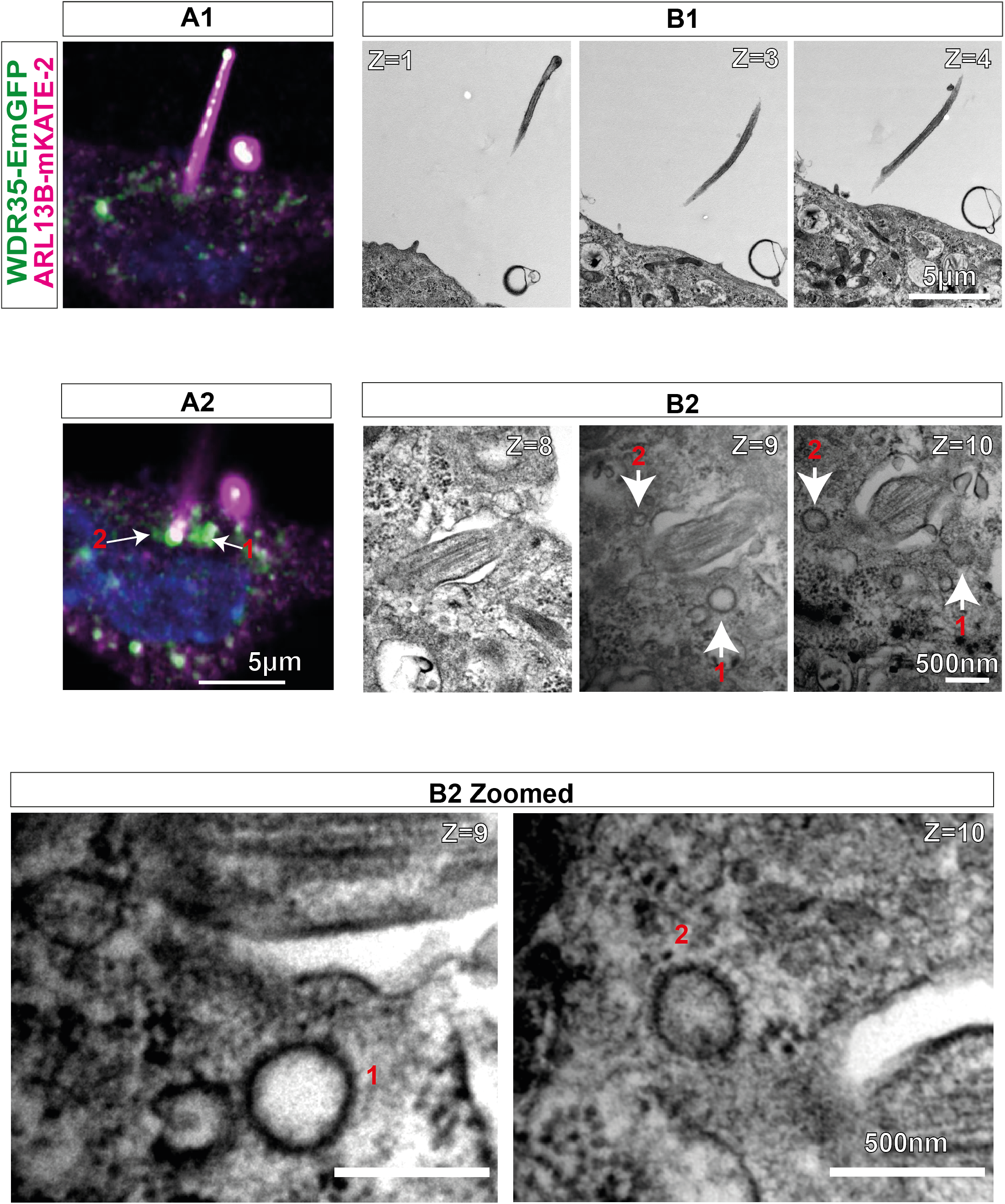
WDR35 is suficient to rescue cilia elongation and restore traficking of coated vesicles, which are GFP-positive by correlative light and electron microscopy. 24 hrs serum-starved *Wdr35*^−/−^ cells rescued for ciliogenesis by expressing WDR35-EmGFP (green) and imaged first with Airyscan confocal imaging followed by TEM imaging. ARL13B-mKATE (magenta) is used as a cilia marker. A1 and A2 represent two sequential Z-stacks from Airyscan confocal imaging. B1 and B2 represent TEM sequential images of 70 nm sections of the same cell. Arrows point at WDR35 localizing close to the cilia base, as shown by LM imaging, and the same corresponds to electron-dense vesicles shown in Z=9 and Z=10 TEM images. The same two sections enlarged in the last panel show two rescued coated vesicles close to cilia. Scale bars: A1, A2 and <u>B1 are 5 µm, and B2 is 500 nm.</u>

**Figure 10.**
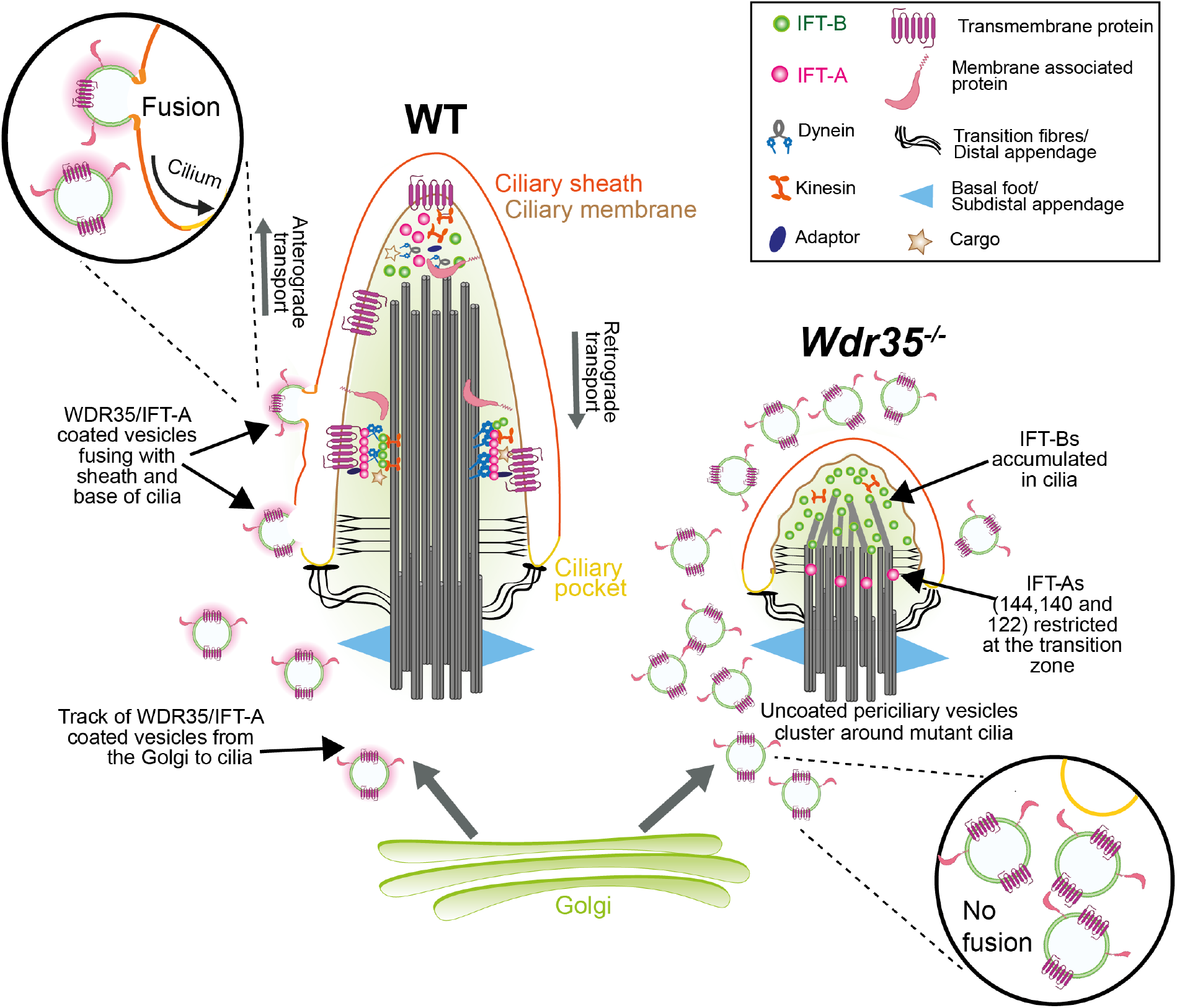
WDR35 and likely other IFT-As assist cargo transport from the Golgi into cilia at the stage of cilia elongation. Diagrammatic representation of the TEM data showing vesicles (blue) with the WDR35 coat (magenta) fusing and localizing around cilia in wild type cells and uncoated vesicles (blue) clustering around cilia in *Wdr35*^−/−^ MEFs. Vesicles follow a track between the Golgi and cilia base in the WT cells but are clustered randomly around cilia in *Wdr35*^−/−^ cells. IFT-B proteins accumulated in short mutant cilia, while IFT-A core components are restricted at the transition zone, and no membrane proteins accumulate inside *Wdr35*^−/−^ cilia, suggesting an arrest at the later stages of ciliogenesis during cilia elongation.

## Discussion

### WDR35 forms a novel coatomer required for Golgi-to-cilia entry of cargo

Vesicle coat proteins, with the archetypal members clathrin and the coat protein complexes I and II (COPI and COPII, respectively), are macromolecular machines that play two central roles in the homeostasis of the cell’s endomembrane system. They enable vesicle formation and select protein and lipid cargo packaged for delivery from a specific donor to functionally segregated compartments. Given the deep sequence and structural similarity of IFT-As to COPI cage proteins, as well as the phenotypic defects in *Wdr35*^−/−^ cells, including lack of ciliary enrichment of a broad range of membrane cargos and the absence of electron-dense cloud on accumulated periciliary vesicles, we propose a novel coatomer function for WDR35 and other IFT-A proteins that is critical for the transport of ciliary membrane cargo into cilia. Two other macromolecular complexes have been proposed to form vesicle associated coats involved in ciliary trafficking: clathrin ***(Kaplan et al., 2010; Molla-Herman et al., 2010)*** and the BBSome complex ***(Jin et al., 2010)***.

Clathrin is a classical coatomer with some documented activity at the ciliary pocket ***(Clement et al., 2013; Pedersen et al., 2016)***. From static images, the directionality of events is difficult to resolve: fission (endocytosis) or fusion (exocytosis). Clathrin vesicles are generally endocytic, where they concentrate cargos and curve off donor membranes for transport into the cytoplasm to fuse with the early endosome compartment. However, a subset of AP-1 clathrin vesicles were shown to traffic between the trans Golgi and basolateral membranes of polarized epithelial cells ***(Fölsch et al., 1999)***. Indeed, in both *C. elegans* ***(Bae et al., 2006; Dwyer et al., 1998; Kaplan et al., 2010; Ou et al., 2007)*** and trypanosomes ***(Vince et al., 2008)*** deletion or depletion of AP-1 leads to defects in cilia assembly and protein trafficking into cilia. However, in mammalian cells, depletion of clathrin and clathrin-associated proteins results in a normal number of cilia with normal lengths ***(Kaplan et al., 2010; Molla-Herman et al., 2010)***, as opposed to the drastically reduced size of *Wdr35*^−/−^ cilia ***(Caparrós-Martín et al., 2015; Fu et al., 2016; Mill et al., 2011)***. This suggests that clathrin is dispensable for vesicular transport into mammalian cilia. Although electron-dense vesicles were observed invaginating from the mammalian ciliary pocket, the electron-density on these vesicular invaginations was unchanged in the absence of clathrin ***(Molla-Herman et al., 2010)***. Using live cell-imaging, the directionality of clathrin-mediated trafficking was reported to be away from cilia ***(Molla-Herman et al., 2010)***. Importantly, we still observe clathrin-coated endocytic structures on the plasma membrane of *Wdr35*^−/−^ cells (**Figure 7 Supplement 2; Movie 6**), and we found no difference in the distribution of clathrin intensity in a volume of 2 µm^3^ around the ciliary base in *Wdr35*^−/−^ cilia compared to controls (**Figure 8 Supplement 1 C-D**).

The BBSome is a macromolecular machine of Bardet-Biedl Syndrome (BBS) proteins which is also postulated to have evolved from an early ancestral coat complex ***(Jékely and Arendt, 2006; van Dam et al., 2013)***. The BBsome shares similar structural elements to the archetypal coatomers and plays a role in cilia function ***(Nachury, 2018)***. In contrast to IFT, mutations in BBSome components including ARL6/BBS3 do not affect cilia assembly and length regulation ***(Domire et al., 2011; Eguether et al., 2014; Lechtreck et al., 2013, 2009; Liew et al., 2014; Nager et al., 2017; Shinde et al., 2020; Xu et al., 2015; Ye et al., 2018)***. Instead, they generally are required for regulating cilia content, mostly for the export of ciliary membrane proteins Although this suggests BBSomes regulate movement of ciliary components between compartments, endogenous localization of BBSome remains unclear, without evidence supporting endomembrane or plasma membrane localization. In contrast, IFT20 has been shown to localize to the Golgi (***Follit et al., 2006; Noda et al., 2016)***. Moreover, whilst there is in vitro evidence that BBSomes can cluster on liposomes, they do not deform membranes, a key step in vesicle formation by coatomers ***(Jin et al., 2010)***. In contrast, purified IFT172, an IFT-B component that is also homologous to COPI *α* and *β*′ like WDR35, can not only assemble on liposomes with high affinity but can also bud 50nm vesicles consistent with coatomer sized products ***(Wang et al., 2018)***. Together this evolutionary conservation of the BBSome with coatomer proteins ***(Jékely and Arendt, 2006; van Dam et al., 2013)***, interaction with in vitro membranes in presence of ARF like GTPase ARL-6, interaction with phospholipids ***(Jin et al., 2010; Nachury et al., 2007)*** and recent cryoEM structures of the complex ***(Chou et al., 2019; Klink et al., 2020; Singh et al., 2020; Yang et al., 2020)*** suggest the BBSome may be working as an adaptor for IFT-A mediated cage formation, similar to other coat adaptors for clathrin (i.e.AP1/AP2) or COP (i.e. *β*-, *γ*-, *δ*-, and *ζ*-COP for COPI). Our data suggests that the electron density observed on vesicles around the ciliary base in control cells is neither clathrin nor BBSome in nature, and is likely composed of WDR35/IFT-A.

### Mechanism of WDR35/IFT-A-assisted coatomer function; regulators of vesicular fusion and fission

Our study demonstrates a requirement for IFT-A to deliver ciliary membrane cargo into cilia, potentially via novel coatomer structure operating between Golgi and ciliary base. Archetypal coatomer protein complexes, including COPII, COPI, and clathrin, concentrate cargo within donor membranes and pinch off vesicles (fission), which then travel to their target organelle membranes, where SNARE and Rab GTPases assist their fusion ***(Bonifacino and Glick, 2004)***. In these cases, the electrondense coats are progressively dismantled such that uncoated vesicles can fuse with acceptor membranes, presumably to facilitate access to the fusion machinery, such as SNAREs, on the surface of the vesicle. As a result of interactions with cargo and lipids with the vesicles, there is evidence that the COPI coat can remain stable on membranes after fission. Moreover, this suggests that COPI vesicle uncoating may be incomplete, such that residual COPI on the vesicle surface enables vesicle recognition and tethering necessary for fusion to the correct acceptor membrane ***(Orci et al., 1998)***. In contrast to the trail of electron-dense vesicles from the Golgi to cilia and a few coated vesicles at the base of cilia in control cilia, we observed ten times more vesicles stalled around the cilia base of *Wdr35*^−/−^ MEFs. These all lack an electron-dense coat suggesting that these transport vesicles are formed but fail to fuse at the ciliary target membrane in the absence of WDR35.

This raises a question as to why a protein like WDR35, which shares structural homology to fission-inducing proteins, gives phenotypes consistent with a fusion-facilitating protein. We speculate that while *Wdr35*^−/−^ MEFs are missing one COPI *α*/*β*′-homolog, the other three core IFT-As (IFT144, IFT140, and IFT122) may be sufficient to compensate by providing interaction motifs necessary for the fission of vesicles from Golgi. Indeed, we show IFT122 to be upregulated in *Wdr35*^−/−^ mutant cells and other IFT-A core proteins are reported to be upregulated in WDR35 patients ***(Duran et al., 2017)***. However, we and others have demonstrated that in the absence of WDR35, the IFT-A complex is unstable ***(Zhu et al., 2017)*** such that any core IFT-A coat on the vesicles from the Golgi may be easily disassembled. It is interesting to note that peripheral IFT139 and IFT43 are helical ***(Taschner et al., 2012)*** similar to SNARE proteins that mediate vesicle fusion with target membranes. Importantly, we show here that these components, which are degraded in the absence of WDR35, could help mediate the fusion of vesicles with the ciliary pocket or base to transfer membrane cargos into the growing cilia sheath. Important next steps will be to systematically investigate vesicular trafficking defects in other IFT-A mutants, as well as identify the GTPase which acts as the energy source to drive fission and fusion of these events.

Recruitment, remodeling, and regulation of protein coats involve cycles of GTP hydrolysis, for example ARF-1 regulates COPI coat ***(Dodonova et al., 2017)***. It is interesting to note that we and others have been unable to purify IFT-A complex with any GTPases ***(Mukhopadhyay et al., 2010)***, suggesting that any interaction is transient. This is even in conditions where we can purify endogenous IFT-B complexes with its associated GTPases IFT22/RABL5 and IFT27/RABL4. In COPI, recruitment of coat components to donor membranes starts with the insertion of small GTPase ARF1 into membranes ***(Dodonova et al., 2017)***. So far only one ARF, ARF4 acting at the TGN ***(Mazelova et al., 2009; Wang et al., 2017)*** has been implicated in ciliary trafficking. However, it plays non-cilial roles, and shows early lethality in mouse knock-outs without affecting cilia assembly ***(Follit et al., 2014)***. Mutations in several related ARLs have defects in cilia structure and/or content, including *ARL3, ARL6* and *ARL13B* ***(Alkanderi et al., 2018; Cantagrel et al., 2008; Fan et al., 2004)***. At least in the case of ARL13B and ARL3, they fail to accumulate and/or enter mutant cilia, even when overexpressed in the absence of WDR35. Other Rab GTPases have been implicated in the ciliary targeting of vesicular cargos ***(Blacque et al., 2018)***. Notably, expression of dominant negative RAB8 in *Xenopus* photoreceptors ***(Moritz et al., 2001)*** results in a strikingly similar accumulation of vesicles to our *Wdr35*^−/−^ mutants which fail to fuse with the ciliary base. Similarly, in RPE-1 cells, dominant negative RAB8 impairs trafficking of ciliary membrane cargos ***(Nachury et al., 2007)***. However, functional redundancy between Rabs may exist as neither single nor *Rab8a;Rab8b* double mutant mice have defects in cilia formation. On the other hand, defects in ciliation were observed when Rab10 was additionally knocked down in *Rab8a;Rab8b* double mutant cells (***Sato et al., 2014)***. Excitingly, our work demonstrates IFT-As to be important for the later stage of ciliogenesis, similar to GTPases like RAB23 ***(Gerondopoulos et al., 2019)*** or RSG-1 ***(Agbu et al., 2018; Toriyama et al., 2016)***. Given that these GTPases have also been shown to sequentially interact with CPLANE subunits INTU and FUZ, which are also required for IFT-A holocomplex assembly ***(Gerondopoulos et al., 2019; Toriyama et al., 2016)***, they will be priorities for future investigations.

Vesicular cargos themselves are also thought to regulate these processes that are necessary to sort and enrich them into appropriate destination vesicles. Several ciliary trafficking motifs have been reported to assist in this process ***(Malicki and Avidor-Reiss, 2014)***. However, our results suggest these may be quite generic motifs as we show mistargeting of a wide range of cilia membrane and membrane associated proteins, in addition to fluorophores carrying lipidated motifs, suggestive of defective trafficking of cilia-bound lipid rafts. Indeed, levels of most endogenous myristoylated proteins and geranylgeranylated proteins, as well as transmembrane proteins, were recently shown by quantitative MS to be strongly reduced in the flagella of *Chlamydomonas* IFT-A mutants ***(Picariello et al., 2019)***. Importantly here, the block in ciliogenesis of an *ift140* null allele was partially overcome by overexpressing a transgene expressing only the TPR domains; this partially rescued cilia length but not ciliary membrane content, further suggesting a full IFT-A coatomer is required for efficient trafficking into cilia.

We have demonstrated that an IFT-A-dependent coat for membrane vesicles derived from the Golgi exists and is necessary for their fusion with the ciliary sheath, which is continuous with the ciliary membrane. We also showed that this coat is necessary, to efficiently deliver cilia-destined signaling molecules into the elongating axoneme of the cilium. Given its efficacy, this IFT-dependent ‘targeted delivery’ module may also be repurposed for other non-ciliary membrane targeting events via polarized exocytosis. Notably in the immune synapse of T cells, the centriole and Golgi apparatus are positioned just below this site of intense vesicle trafficking ***(Finetti et al., 2009)***. In these cells with ‘frustrated centrosomes’, IFT20 is required for rapid clustering of TCRs necessary for T cell activation ***(Finetti et al., 2009)***. Work in the neuroretina with immunogold TEM revealed IFT localization to vesicles tracking towards the postsynaptic membranes of secondary retinal cell dendrites, found at great distance from the connecting cilium of photoreceptors ***(Sedmak and Wolfrum, 2010)***. Dendritic exocytosis is implicated in retrograde signaling, synaptic plasticity, and neuronal morphology ***(Kennedy and Ehlers, 2011)***. Future studies into this IFT-dependent coatomer and the membrane trafficking processes it controls may expand our phenotypic understanding of the ciliopathies beyond the cilium.

### Methods and Materials

Preparation of primary MEFs, cell culture, ciliation and genotyping Primary MEFs were harvested from E11.5 embryos and cultured in complete media [Opti MEM-I (Gibco, REF: 31985-047) supplemented with 10% foetal calf serum (FCS) and 1% penicillin-streptomycin (P/S) and 0.026*µ*l *β*-mercaptoethanol] and incubated at 37°C in a hypoxic incubator (3% O_2_ and 5% CO_2_). To induce ciliogenesis, 70-80% confluent cells were serum-starved for 24 hr. Genotyping was done as described before for the *Wdr35* line ***(Mill et al., 2011)*** and *Dync2h1* line ***(Caparrós-Martín et al., 2015)***. *Pcm1-SNAP* mouse line was made by Dr. Emma Hall (Hall E. et al., unpublished) by endogenous tagging of *Pcm1* by CRISPR. *Pcm1SNAP* mouse line was crossed with *Wdr35*^+/−^ and genotyped to screen E11.5 embryos homozygous for both *Wdr35*^−/−^ and *Pcm1*^*SNAP* /*SNAP*^. MEFs prepared from these embryos were used to image PCM-1 localization in WT and *Wdr35*^−/−^ using antibodies and other reagents listed in **Table 1**.

**Table 1.** List of secondary antibodies.

| Antibody | Host | Source | Dilution | Application |
| --- | --- | --- | --- | --- |
| ECL α-Mouse IgG, HRP-conj | Sheep | GE Healthcare NA931-1ML | 1:10,000 | WB |
| ECL α-Rabbit IgG, HRP-conj | Goat | GE Healthcare RPN4301 | 1:10,000 | WB |
| α-Rabbit Light-Chain specific HRP conj | Mouse | Merk-MAB201P | 1:10,000 | WB |
| α-Rabbit IgG Light-Chain Specific mAb | Mouse | CST-L57A3 | 1:10,000 | WB |
| Alexa 488, 594, 647 conj-α-Mouse | Donkey | Molecular Probes | 1:500 | IF |
| Alexa 488, 594, 647 conj-α-Rabbit | Donkey | Molecular Probes | 1:500 | IF |
| Alexa 488, 594, 647 conj-α-Goat | Donkey | Molecular Probes | 1:500 | IF |

### Electroporation of MEFs

Cells were trypsinized to a single-cell suspension and resuspended in 10 *µ*l Resuspension Buffer R per 0.5×10^5^*cells*/*transfectionreaction, mixedwithplasmidDNA*(0.75*µ*g/transfection) **Table 3** and electroporated (voltage 1350 V, width 30 ms, one pulse) using a Neon Nucleofection kit (ThermFisher MPK-1096), according to the manufacturer’s protocol. Transfected cells are harvested or visualized 24-48 hr post electroporation.

| Antibody | Species | Source | Dilution | Fixation |
| --- | --- | --- | --- | --- |
| ARL13B | Rabbit | Proteintech 17711-1-AP | 1:100 IF | PFA |
| Ac α-tubulin | Mouse | Sigma T6793 | 1:1000 IF | PFA/GA |
| Ac α-tubulin | Rabbit | Abcam ab179484 | 1:1000 IF | PFA/GA |
| γtubulin (GTU88) | Mouse | Sigma T6557 | 1:100 IF | PFA |
| GFP | Mouse | Roche (11814460001) | 1:1000 IF 1:1000 WB | PFA |
| GFP | Rabbit | Abcam (ab6556) | 1:1000 IF 1:1000 WB | PFA |
| GFP | Rabbit | Santa Cruz SC8334 | 1:5000 WB | - |
| IFT88 | Rabbit | Proteintech 13967-1AP | 1:200 IF 1:1000 WB 3μg IP | PFA |
| IFT81 | Rabbit | Proteintech 11744-1-AP | 1:200 IF 1:1000 WB | PFA |
| IFT144 | Rabbit | Proteintech 1367-1-AP | 1:200 IF 1:1000WB | PFA |
| IFT140 | Rabbit | Proteintech 17460-1-AP | 1:200 IF 1:1000 WB 3μg IP | PFA |
| IFT122 | Rabbit | Proteintech 19304-1-AP | 1:200 IF 1:1000 WB | PFA |
| IFT122 | Rabbit | Aviva ARP 53817-P050 | Not for IF 1:1000 WB | - |
| IFT43 | Rabbit | From Victor Ruiz-Perez, UAM | 1:200 IF 1:2000 WB | PFA |
| IFT139 | Rabbit | Novus-NBP1-90416 | 1:1000 WB 1:200 IF | PFA |
| IFT121 | Rabbit | Custom made Proteintech | 1:50 WB Not for IF | - |
| PCM-1 | Rabbit | Proteintech 19856-1-AP | 1:100 IF | MeOH |
| SNAP | Rabbit | New England Biolabs- P9310S | 1:300 IF | MeOH |
| MKS-1 | Rabbit | Proteintech 16206-1-AP | 1:100 IF | MeOH |
| NPHP-1 640 | Rabbit | from Greg Pazour, UMass | 1:100 IF | MeOH |
| ARL3 | Rabbit | Proteintech 10961-1-AP | 1:100 IF | PFA |
| SiR-Tubulin | - | Spirochrome (SC002) | 100nM | Live |
| SNAP-TMR | - | SNAP-CellTMR-STAR (S9105S) | 1μM | Live |

### Endogenous IFT IPs

Embryos were lysed and homogenized in IP lysis buffer (10*µ*l/mg) at 4°C on a rotator for 30 minutes. Composition of IP lysis buffer is [(50mM Tris-HCl (pH 7.5), 100mM NaCl, 10% Glycerol, 0.5mM EDTA, 0.5% IGEPAL, and 1/100 Halt protease and phosphatase inhibitor (ThermoScientific: REF78443) and a tablet of protease inhibitor tablet - 1 tablet per 10mL (Complete Mini, Roche-REF 11836170001). The lysate was cleared by spinning at 4°C, 14,000rpm, for 20 minutes. The protein concentration was determined using the BCA Protein Assay Kit as per manufactures instruction (ThermoFisher REF 23225). For each IP, 500µg of protein was incubated with 3*µ*g of each antibody overnight at 4°C **Table 1** with mild agitation (side-to-side). Immunoprecipitation of immunocomplexes was done using PureProteome™ Protein G magnetic beads (Millipore LSKMAGG10). 30*µ*l beads/IP were equilibrated with 500*µ*l IP lysis buffer by gentle agitation for 5 min at 4°C. Tubes were placed on a magnet for 2min, and the buffer was aspirated off with the fine pipette. 200µl antibody-lysate mix was added to each tube of 30µl equilibrated beads and incubated for 45 min with agitation, to concentrate immunoglobulin complexes on beads at 4°C. Washes (8 times) were performed, each lasting 5min. Washes were as follows: 2x washes in Wash Buffer-1 (same as IP lysis buffer), followed by 2x washes with Wash Buffer-2 (IP lysis buffer with reduced 0.2% IGEPAL), finally 4x washes with Wash Buffer-3 (IP lysis buffer without any IGEPAL detergent). All wash buffer is aspirated, and dry beads were stored at −80°C, or samples were sent immediately for mass spec.

### Mass spectrometry

All mass spectrometry experiments were done at the IGMM Mass Spectrometry facility, as per their published protocol ***(Turriziani et al., 2014)***. Briefly, the immunocomplexes collected on magnetic beads were processed to generate tryptic peptides. Proteins were eluted from beads by incubating at 27°C for 30 min in elution buffer (2M urea, 50mM Tris-HCl pH 7.5 and 5*µ*g/mL trypsin). The sample was centrifuged, bead pellets washed twice and the supernatant from samples digested overnight at room temperature. Iodoacetamide was added to the samples to inhibit disulfide bond formation and incubated for 30 min in the dark. Followed this, trifluoroacetic acid (TFA) was added to stop tryptic digestion. Desalting and pre-fractionation of the digested peptides were done by manually using C18 pipette stage-tips filled with 3M empore disc activated with 50% acetonitrile and 0.1% TFA and then washed once with 0.1% TFA. The peptide mixtures were passed manually along to the column with a syringe to concentrate and purify the analytes. Peptides were subsequently eluted twice in 50% acetonitrile and 0.1% TFA and both eluates were combined. Samples were concentrated and resuspended in 0.1% TFA. This was followed by chromatographic separation on a Reprosil column along a 3-32% acetonitrile gradient. The LC setup was attached to a Q-Exactive mass spectrometer, and ion mass spectra were obtained following HPLC during a tandem MS run. Mass spectra were analyzed using MaxQuant software. Label-free quantification intensity (LFQ) values were obtained for analysis by identifying mass/charge ratio, and their intensities at a specific elution time for individual peptides. The data was collected for both control (GFP) and specific proteins IPs (i.e., IFT88, IFT140 - **Table 1**). LFQ values for the proteins were obtained by summing the ion intensities corresponding to peptides after assigning the unique peptides to proteins. The ratio of LFQ intensities of test: control was taken, where higher the ratio better corresponds to a better enrichment of protein in complex. Complete Mass spec data is available on ProteomeXchange **(identifier PXD022652)**. The relative concentration of IFTs was calculated after normalizing the individual test values with respective GFP-LFQs, as shown in the figures.

### Western Blots

Cells or tissues were lysed in 1X cell lysis buffer with the addition of 1/100 Halt protease and phosphatase inhibitor (Thermo Scientific: REF78443) and a tablet of protease inhibitor tablet, 1 tablet per 10mL (Complete Mini, Roche-REF 11836170001). 1X cell lysis buffer is (Cell Signalling Technology: 20mM Tris-HCl (pH 7.5), 150mM NaCl, 1mM Na_2_ EDTA, 1mM EGTA, 1% Triton-X100, 2.5Mm sodium pyrophosphate, 1mM *β*-glycerophosphate, 1mM Na3VO4, 1µg/ml leupeptin). The lysate from embryos was homogenized at 4°C for 30 min and from cells was sonicated briefly (5x,10-second pulses, Bioruptor Diagenode) to lyse the tissue or cells. The lysate was centrifuged at 14,000g at 4°C for 30 min and the supernatant transferred to a fresh tube. Ready-to-use SDS-PAGE gels (NuPage Novex precast gels, ThermoFisher) were used to separate proteins. The resolved proteins on the gel were transferred to PVDF (Hybond P, GE HealthCare) using the XCell II Blot module as per manufacturer’s instruction. The membrane was then blocked with a 10% solution of dried skimmed milk (Marvel Premier Foods) made in 1X TBST (0.05% Tween-20 in TBS) for 1 hr RT, washed with PBS and incubated with primary antibody **Table 1** diluted in 1% skimmed milk solution in 1X TBST overnight at 4°C on shaker/roller. Membranes were washed in 1X TBST 3X, 10min followed by a 1X wash with PBS, and incubated in HRP-conjugated secondary antibody from appropriate species **(Table 2)** for one hour at RT, diluted in a solution of 1X TBST and 1% milk. Blot was then washed with 1X TBST, three times and with PBS twice. After the washes, protein bands were detected by the Super Signal ELISA Femto kit (ThermoScientific: 37074) or Super Signal ELISA Pico kit (Thermo Fisher: 37069). Protein bands were visualized digitally by transmission light imaging on ImageQuant LAS 4000 (GE HealthCare) and analyzed using ImageQuant TL software. Protein bands on blots were quantified with ImageJ/FIJI software by measuring individual bands intensity and normalizing intensities with loading control bands on the same blot.

**Table 2.**
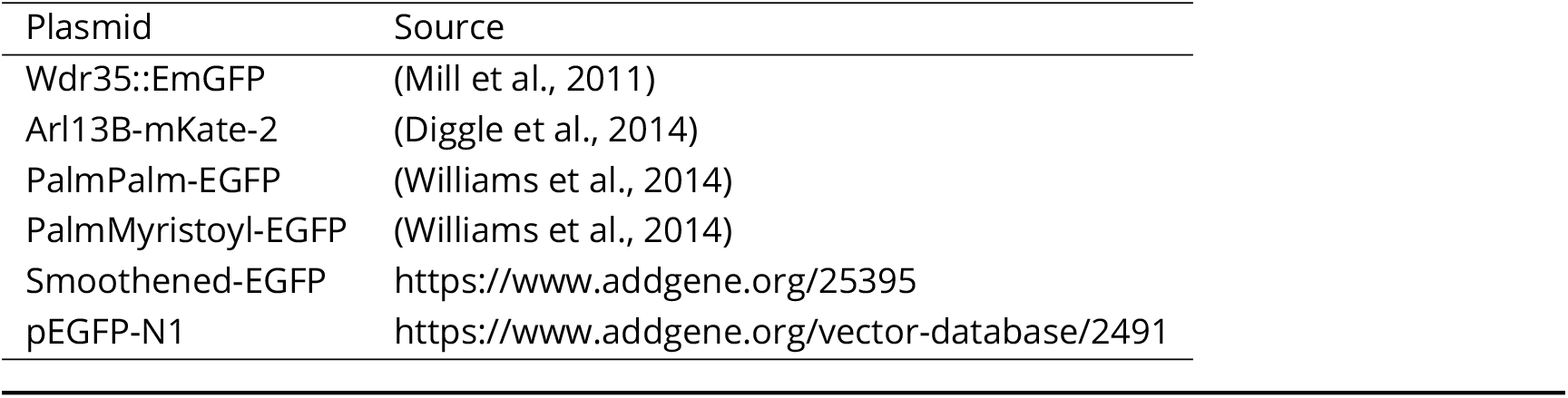
List of plasmids

| Plasmid | Source |
| --- | --- |
| Wdr35::EmGFP | (Mill et al., 2011) |
| Arl13B-mKate-2 | (Diggle et al., 2014) |
| PalmPalm-EGFP | (Williams et al., 2014) |
| PalmMyristoyl-EGFP | (Williams et al., 2014) |
| Smoothened-EGFP | https://www.addgene.org/25395 |
| pEGFP-N1 | https://www.addgene.org/vector-database/2491 |

### Immunofluorescence

Cells were washed two times with warm PBS, then fixed in either 4% PFA in 1X PHEM/PBS 15 min at room temperature/ 2% fresh glutaraldehyde in 1X PHEM for 15 min, or 100% cold methanol for 3 min according to the Ab antigenicity compatibility **Table 1**, washed twice with PBS. 1X PHEM (pH 6.9) contains: 60mM PIPES, 25mM HEPES, 10mM EGTA, and 4mM MgSO_4_·7H_2_0). The cells were treated with 50mM NH_3_Cl, two times, 15 min each for PFA fixed cells, or 0.01mg of NaBH_4_ in 1X PBS for 7 min for glutaraldehyde fixed cells to quench autofluorescence. Cells were then washed twice with PBS. Cells were permeabilized with 0.25% Triton-X 100 in 1X TBS for 10 min at room temperature. Cells were rinsed twice in 1X TBS for 5 min. Blocking for non-specific binding was done by incubating samples in 10% donkey serum in 1X TBS and 0.2% Tween-20 for 60 min at room temperature. Samples were washed twice with PBS. Primary antibodies **(Table 1)** were added to samples and incubated for 60 min at room temperature or 4°C overnight, in dilutant made of 1% donkey serum in 1X TBS and 0.025% Triton X-100. Samples were washed with 0.25% Triton-X 100 in TBS 4-6 times 10 min each. Secondary antibodies diluted in 1% donkey serum and 0.025% Triton-X100 in 1X TBS were incubated on samples for 60 min room temperature. Samples were washed with TBST 4-6 times 10 min, stained with DAPI (1:1000) in PBS for 5 min at room temperature, again washed with PBS and directly imaged or coverslips were added on slides using ProLong Gold antifade (Thermo Fisher Scientific), according to the manufacturer’s instructions. Confocal imaging for both fixed and live-cell samples was done on Leica SP5 using the LAS-AF software, 405 nm diode, Argon and 561 and 648 nm laser lines, three Photomultiplier tubes, and one HyD GaSP detector as per the requirement of the experiment. Images were scanned using a 63X 1.4NA oil immersion objective and latter processed using ImageJ and Imaris software.

### IFT-A sequence homology search and structural modeling

The sequence match of IFT-A proteins was found by iterative rounds of homology searches via alignment for sequence proximity-based clustering as described before ***(Wells et al., 2017; Wells and Marsh, 2019)***. Further Swiss Model server was used to model IFT-A complex protein structures as described on the server ***(Waterhouse et al., 2018)***. Briefly, a template search with BLAST and HHblits was performed against the SWISS-MODEL template library. The target sequence was searched with BLAST against the primary amino acid sequence contained in the SMTL. An initial HHblits profile, followed by one iteration of HHblits against NR20, was run and the obtained profile then searched against all profiles of the SMTL. The top hit in all of IFTA searches was 3mkqA ***(Lee and Goldberg, 2010)***, a coatomer *β*′ subunit 2.5Å X-ray structure with 14% - 20% sequence identity and 25% - 30% sequence similarity with different IFT-A proteins. A coatomer *α* subunit was also found within these top matches. Models were built on the target-template alignment using ProMod3. Coordinates that are conserved between the target and the template were copied from the template to the model. Insertions and deletions were remodelled using a fragment library. Side chains were then rebuilt. Finally, the geometry of the resulting model was regularized by using a force field. In case loop modeling with ProMod3 fails, an alternative model was built with PROMOD-II. The global and per-residue model quality has been assessed using the QMEAN scoring function. The obtained model was processed later in Pymol software for structural analysis.

### Transmission electron microscopy

TEM sample preparation: 24 hr serum-starved MEFs were chemically fixed for flat embedding using the following protocol: (1) Cells were grown on 60mm dishes, and ciliogenesis was induced by serum starvation for 24 hr. (2) For prefixation under culture conditions, 25% glutaraldehyde was added to the growth medium to a final concentration of 1%, mixed gently, and incubated for a few minutes at 37°C. (3) The growth medium (containing the glutaraldehyde) was replaced with a sample buffer (0.1M HEPES, 4mM CaCl_2_, pH 7.2) containing 2% glutaraldehyde and incubated 1 hr at room temperature (replacing the fixation buffer with fresh one after 20 min). All prefixation solutions were pre-warmed to 37°C, and all steps were done at 37°C, to preserve the cytoskeleton. (4) The fixation buffer was replaced with fresh fixation buffer and incubated for 4hrs at 4°C. (5) After that, the sample was washed once in sample buffer and 2–3 times in distilled water, each for 5–10 min, gently removing and replacing the buffer. (6) Samples were incubated in 1% OsO_4_ (EMS) (in distilled water) for 1 hr at 4°C, (7) washed 3–4 times for 10 min each in distilled water, and (8) incubated in 1% uranyl acetate (EMS) in distilled water overnight at 4°C. (9) Then, samples were rinsed 3–4 times for 10 min each in distilled water and (10) dehydrated using a graduated series of ethanol: 30%, 50%, 70%, 80%, 90%, 96% ethanol, 5 min each step at 4°C, followed by twice rinsed in anhydrous 100% ethanol 10 min each at RT. (11) Infiltration was performed using a 1:1 mixture of LX112 (Ladd Research, USA; EMS) and ethanol 2 hr, followed by pure LX112 overnight and another 2 hr pure LX112, where all steps were performed at room temperature. (12) Flat embedding: For flat embedding, the caps of the BEEM embedding capsule (size, No3, EMS) were cut off and capsules filled with LX112. The capsules were inverted over a selected area of the cell monolayer in the dish, and the resin cured at 60°C oven for 48 hr. The capsule was then removed by breaking off from the dish, leaving the monolayer cells embedded in the surface of the block. The top 1-2 mm block surface with embedded cell monolayer was removed by sawing it off and remounting it on a dummy block for sectioning in a perpendicular orientation with respect to the monolayer. (13) Sectioning and post-staining: For sectioning and post-staining, 300 nm thick serial sections were cut by Leica Ultracut UCT (Leica microsystem, Wetzlar, Germany) with a diamond knife and sections picked up with a Formvar (EMS) coated 1×2 mm slot copper grid (EMS). Sections were post-stained with 2% uranyl acetate for 10 min, then with lead citrate for 5 min. Imaging: Sections were stained on the grid with fiducials (15 nm gold nanoparticles, Sigma-Aldrich). 70 nm thick sections were cut for regular TEM imaging, and 300 nm thick sections were prepared for tomographic acquisition. Tilt series were acquired on a Tecnai F30 (FEI) transmission electron microscope, operated at 300 kV, and equipped with 2048×2048 Gatan CCD camera. The SerialEM software ***(Mastronarde, 2005)*** was used for automatic acquisition of double tilt series. Tomographic tilt series were recorded with a pixel size of 1.235 nm, a maximum tilt range of about 60°, and tilt steps of 1°. Tomographic reconstruction, joining of tomograms from consecutive sections, segmentation, and visualization of the tomograms was done using the IMOD software package ***(Kremer et al., 1996)***. 24hr serum-starved WT, *Wdr35*^−/−^, and *Dync2h1*^−/−^ cells were serially sectioned parallel to the adherent surface. Two to four 300nm parallel serial sections are required to get the whole 3D volume ultrastructural view covering full cilia and their cellular surroundings. We reconstructed 45 tomograms to get a minimum of 3-4 whole cell volumes for each genotype. We took micrographs of 30 WT, 20 *Wdr35*^−/−^, and 30 *Dync2h1*^−/−^ cells for this study.

### CLEM (correlative light and electron microscopy)

WDR35-EmGFP and ARL13B-mKate expressing *Wdr35*^−/−^ MEFs were serum-starved for 24 hr, stained with Hoechst 33342 (R37605) for 10 min in culture condition, fixed with 4% PFA and 0.1% GA in 1X PHEM and imaged on Zeiss LSM 880 upright single photon point scanning confocal system with Quasar detector (32 spectral detection channels in the GaAsP detector plus 2PMTs) and transmitted light detector, Airy scan detector for high-resolution imaging. Cells were grown on 35mm glass bottom dishes with grids (Cat. No. P35G-1.5-1.4-C-GRID) and firstly brightfield images were made with Plan-Apochromat 10X/0.45 M27 objective to save the coordinates of cells needed for the correlation with the respective TEM data. Confocal and airy scan imaging was done using Plan-Apochromat 63x/1.4 oil DMC M27 objective, 405 nm laser diode, 458, 477, 488, 514 nm multiline integrated Argon laser and 594 nm integrated HeNe laser. Z-stack was acquired sequentially to get the whole 3D volume of the cell and the image was further deconvolved using the inbuilt software.

After Airy scan imaging the sample was processed for TEM as described above. 70 nm sections were made for the regions of saved coordinates from brightfield imaging, mounted on grids and imaged on FEI Morgagni TEM (100kV) microscope.

### Image analysis and measurements

All image processing and statistics was done on FIJI (http://fiji.sc/Fiji). Macros for PCM-1 (RadialIn-tensityFromCentrosomes.ijm) and clathrin (3DMeanIntensityfromUserDirectedPoints.ijm) quantifi-cation can be found on GitHub (https://github.com/IGMM-ImagingFacility/Quidwai2020 WDR35paper). To measure PCM1 intensity radially from the centrosomes, an average intensity projection of the z-stack was obtained, then the gamma-tubulin signal was segmented using RenyiEntropy threshold and the Analyze Particles tool to obtain the centrosomes. The selections obtained were enlarged using the “Make Band” function to create a band. This was done by increasing in 1 µm increments until there were five bands. The centrosome masks and the surrounding bands were measured on the PCM1 channel of the average intensity projection image. To quantify clathrin intensity around the cilia base, a point was selected as the center of the basal point. The user blinded to file name and condition while quantification took place. This point was expanded 1 µm in each direction to create a shall of 2 µm diameter in x,y, and z ***(Ollion et al., 2013)***. This shell was then measured using the 3D image suite in ImageJ. IMOD and ETOMO (https://bio3d.colorado.edu/imod/) was used to reconstruct tomograms and segmentation of tomograms respectively.

### Statistics

All statistical analysis was carried out using GraphPad Prism 8 (version 8.4.1; GraphPad software, USA) as described in the text.

## Acknowledgments

We thank the IGMM Advanced Imaging Resource and Mass Spectrometry facility (in particular Jimi Wills and Alexander von Kriegsheim). We thank the Electron Microscopy Facility (in particular Tobias Fürstenhaupt, Michaela Wilsch-Bräuniger and Daniela Vorkel) and the Light Microscope Facility from the Services and Facilities of the Max Planck Institute of Molecular Cell Biology and Genetics, Dresden (in particular Sebastian Bundschuh). We thank Esben Lorentzen, Toby Hurd, Ian Jackson and Patricia Yeyati for helpful discussions and comments on the manuscript.We are grateful to Greg Pazour (UMass) and Victor Ruiz (UAM) for sharing custom antibodies, and Jonathan Eggen-schwiler (UGA) for Ift122::GFP used in this study. This work was supported by a Ph.D. studentship to T.Q., funded through a grant from the UK Medical Research Council to the Edinburgh Super Resolution Imaging Consortium (ESRIC) and an EMBO Short Term Fellowship (No. 7961). J.A.M. is a Lister Institute Research Prize Fellow. Work in the group of G.P. and in the MPI facilities was supported by the Max Planck Society and by the European Research Council (ERC) under the European Union’s Horizon 2020 research and innovation programme (grant agreement No. 819826) to G.P. Work in the group of P.M. is supported by MRC core funding (MC_*U*_ *U*_1_2018/26).

## Author contributions

T.Q. developed the project, performed the bulk of the experiments (including FM, TEM, tomography, and segmentation, CLEM, protein work, homology modeling and cell culture), quantified and analysed the data, interpreted results and prepared figures. E.H generated a mouse line with SNAP tagged PCM-1 (unpublished work), contributed to its FM and quantification. L. M. wrote scripts for clathrin and PCM-1 quantification. M.A.K maintained mouse lines. J.N.W and J.A.M. contributed to bioinformatic analysis. W.L. prepared samples for EM imaging. P.K. contributed to CLEM data. G.P contributed to data analysis and results interpretation, supported and provided resources funding for the EM and CLEM part of the project. P.M. conceived and supervised the project, contributed to data, its analysis and results interpretation, as well as provided resources and funding. T.Q. and

P.M. wrote the manuscript, and all authors contributed to its editing.

## Movie Legends

**Movie 1. The organization of centriolar satellites is not disrupted in *Wdr35***^−/−^ **mutants**. *Wdr35*^+/+^;*Pcm1*^*SNAP* /*SNAP*^ and *Wdr35*^−/−^; *Pcm1*^*SNAP* /*SNAP*^ MEFs electroporated with ARL13B-EGFP (green), serum-starved for 24 hrs andstained for SiR-tubulin (grey) and SNAP-TMR (magenta) and imaged live on LEICA SP5 microscope using a 63X, 1.4 oil immersion objective. The movie is compiled as 5 fps. PCM-1 density around cilia is not altered in the absence of WDR35. **(Related to Figure 1 Supplement 2C)**.

**Movie 2. Cilia specific membrane-associated cargo (A) ARL13B, and membrane-integrated cargo (B) SMO fail to localize in *Wdr35***^−/−^ **cilia**. WT and *Wdr35*^−/−^ MEFs expressing ARL13B-EGFP (green) and Smoothened-EGFP (green), serum-starved for 24 hrs, stained with SiR-tubulin and imaged live on LEICA SP5 microscope using a 63X, 1.4 oil immersion objective. The movie is compiled as 5 fps. **(Related to Figure 5)**.

**Movie 3. Track of electron-dense vesicles are present between Golgi and cilia in control mouse fibroblast**. 24 hr serum-starved cells are prepared for EM analysis by plastic embedding and making 300 nm thick sections. Tomogram reconstructed from two 300 nm sections stitched together shows the presence of electron-dense vesicles between the Golgi and cilia. The 3D volume shown in the upper half is segmented in the lower half of the movie. Daughter centriole (blue), basal body (purple), ciliary membrane (brown), ciliary sheath (orange), ciliary pocket (yellow), Golgi (green), and vesicles with dense electron clouds are shown in magenta. Arrows are pointing at the track of vesicles between the Golgi and cilia. **(Related to Figure 7A)**.

**Movie 4. Electron-dense vesicles are observed around the base of cilia in control mouse fibroblasts**. 24 hr serum-starved cells are prepared for EM analysis by plastic embedding and making 30 0nm thick sections. Tomogram reconstructed from the 30 0nm thick cell section shows electron-dense vesicles clustering at the base of cilia. The 3D volume shown in the upper half is segmented in the lower half of the movie. The basal body (purple), ciliary membrane (brown), ciliary sheath (orange), ciliary pocket (yellow), Golgi (green), and vesicles with dense electron clouds are shown in magenta. **(Related to Figure 7B)**.

**Movie 5. In *Wdr35***^−/−^ **fibroblasts, an accumulation of small uncoated vesicles are present around short and stumpy cilia**. 24h r serum-starved cells are prepared for EM analysis by plastic embedding and making 300 nm thick sections. Tomogram reconstructed from tilt series of two 300 nm thick cell sections and stitched together shows ten times more vesicles randomly clustering around cilia in the mutant. The 3D volume shown in the upper half is segmented in the lower half of the movie. The basal body (purple), ciliary membrane (brown), ciliary sheath (orange), ciliary pocket (yellow), Golgi (green), and uncoated vesicles around the cilia is shown in cyan. The ciliary membrane is floppy compared to WT MEFs and axonemal microtubules are poorly polymerized. **(Related to Figure 7C)**.

**Movie 6. In *Wdr35***^−/−^ **fibroblasts, preciliary vesicles fail to fuse with ciliary pocket or ciliary sheath**. 24 hr serum-starved cells are prepared for EM analysis by plastic embedding and making 300 nm thick sections. Tomogram reconstructed from tilt series of three 300 nm thick cell sections and stitched together shows the transition zone is unaltered in the mutant. Vesicles lacking any electron dense cloud cluster around mutant cilia, but fail to fuse with ciliary sheath or ciliary pocket. The 3D volume shown in the upper half is segmented in the lower half of the movie. Daughter centriole (dark blue), basal body (purple), ciliary membrane (brown), ciliary sheath (orange), ciliary pocket (yellow), basal foot/ subdistal appendage (red), transition fibers/ distal appendage (orchid), Y-Links (white), Golgi (green), and vesicles lacking any electron dense cloud present around cilia (cyan). Arrows are pointing at the clathrin-coated vesicles budding from the cell plasma membrane. **(Related to Figure 7 Supplement 2)**.

**Movie 7. *Dync2h1***^−/−^ **cilia lack both coated as well as uncoated vesicles at the cilia base, whilst ectosomes are seen budding from the tip**. 24 hr serum-starved *Dync2h1*^−/−^ MEFs are prepared for EM analysis by plastic embedding and making 300 nm thick sections. Tomogram reconstructed from the tilt series of three 300 nm sections and stitched together shows a sturdy ciliary membrane, well-polymerized microtubules in the axoneme, almost no coated or uncoated vesicles at the cilia base, and ectosome vesicles could be seen budding from the tip of cilia. A 40 nm striped pattern could be seen present throughout the length of cilia. Arrows point at the basal body (purple), cilia (magenta), and ectosome (green). This is also an example of the rare event of having two cilia in the same ciliary sheath. **(Related to Figure 7 Supplement 3C-Cell1)**.

---

**Figure 1–Figure supplement 1.**
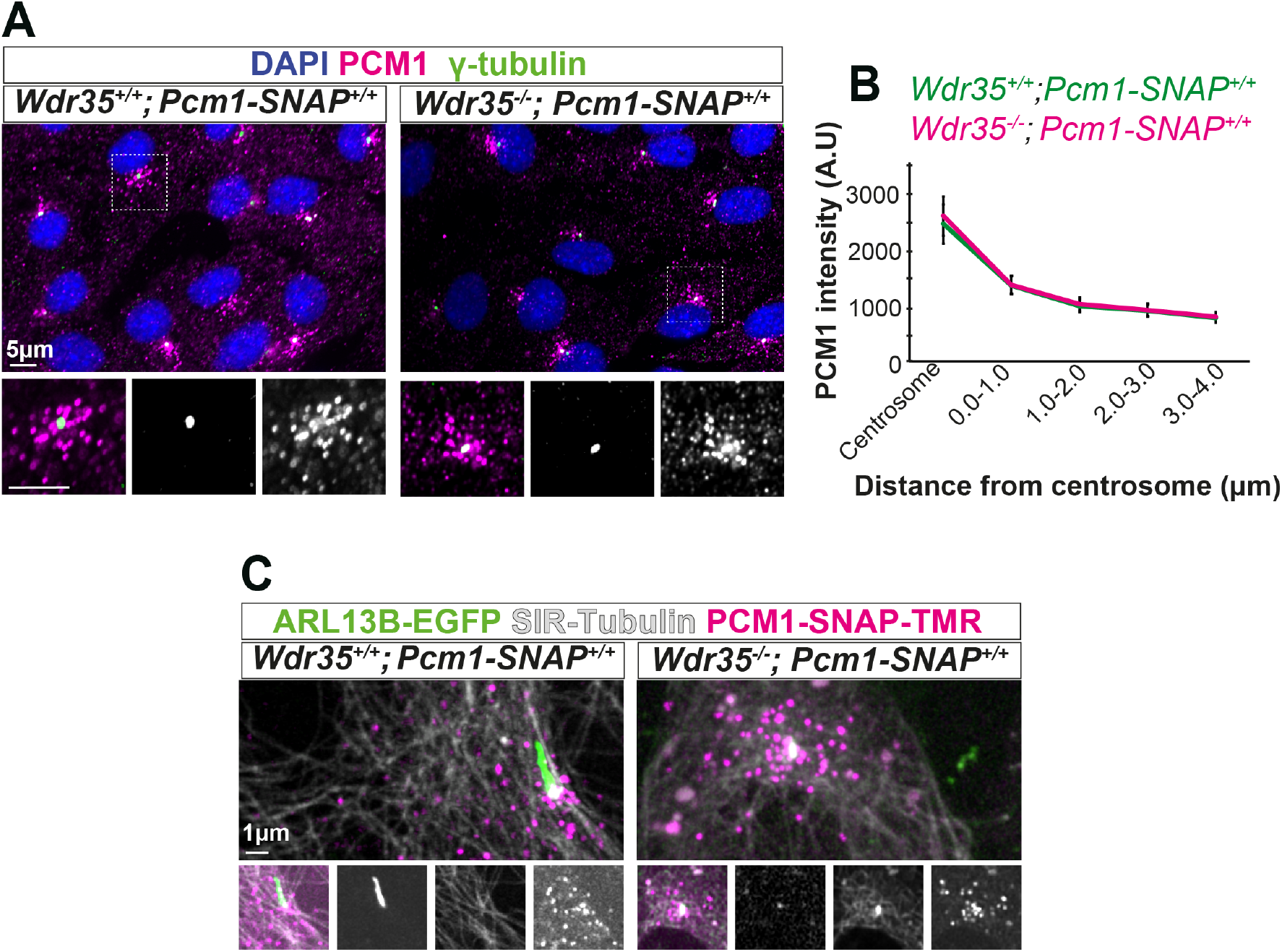
The organisation of centriolar satellites (CS) around Wdr35 ^−/−^ cilia is not changed. CS marker PCM1 intensity and localisation are unchanged in *Wdr35*^+/+^and *Wdr35*^−/−^ MEFs serum starved for 24h to induce ciliogenesis and imaged (A) fixed after staining with antibodies; PCM1 (magenta) and *γ*-tubulin (green). Nuclei are in blue. (C) Imaged live after staining with SNAP-TMR dye for endogenous SNAP tagged PCM1 (magenta) and microtubule marker SiR-tubulin (grey). These cells are also expressing ARL13B-EGFP (green) (**Movie 1)**. (B) Quantification of PCM1 intensity around the centrosome in concentric rings of 1 *µ*m around the basal body. n=50 cells (3 biological replicates each).

**Figure 6–Figure supplement 1.**
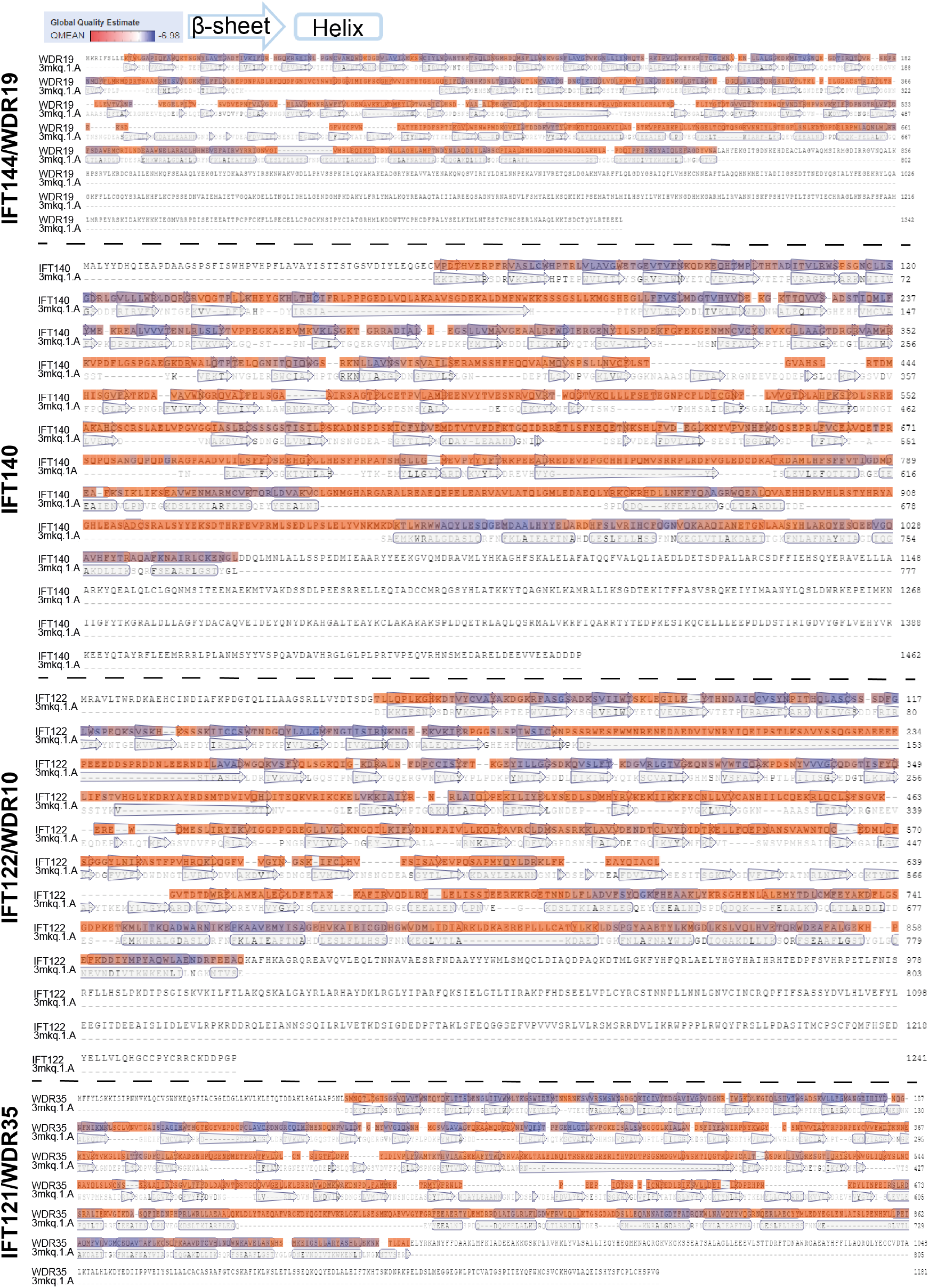
Sequence blast result between IFT-A proteins and COPI-/’ received from SWISS MODEL server.

**Figure 7–Figure supplement 1.**
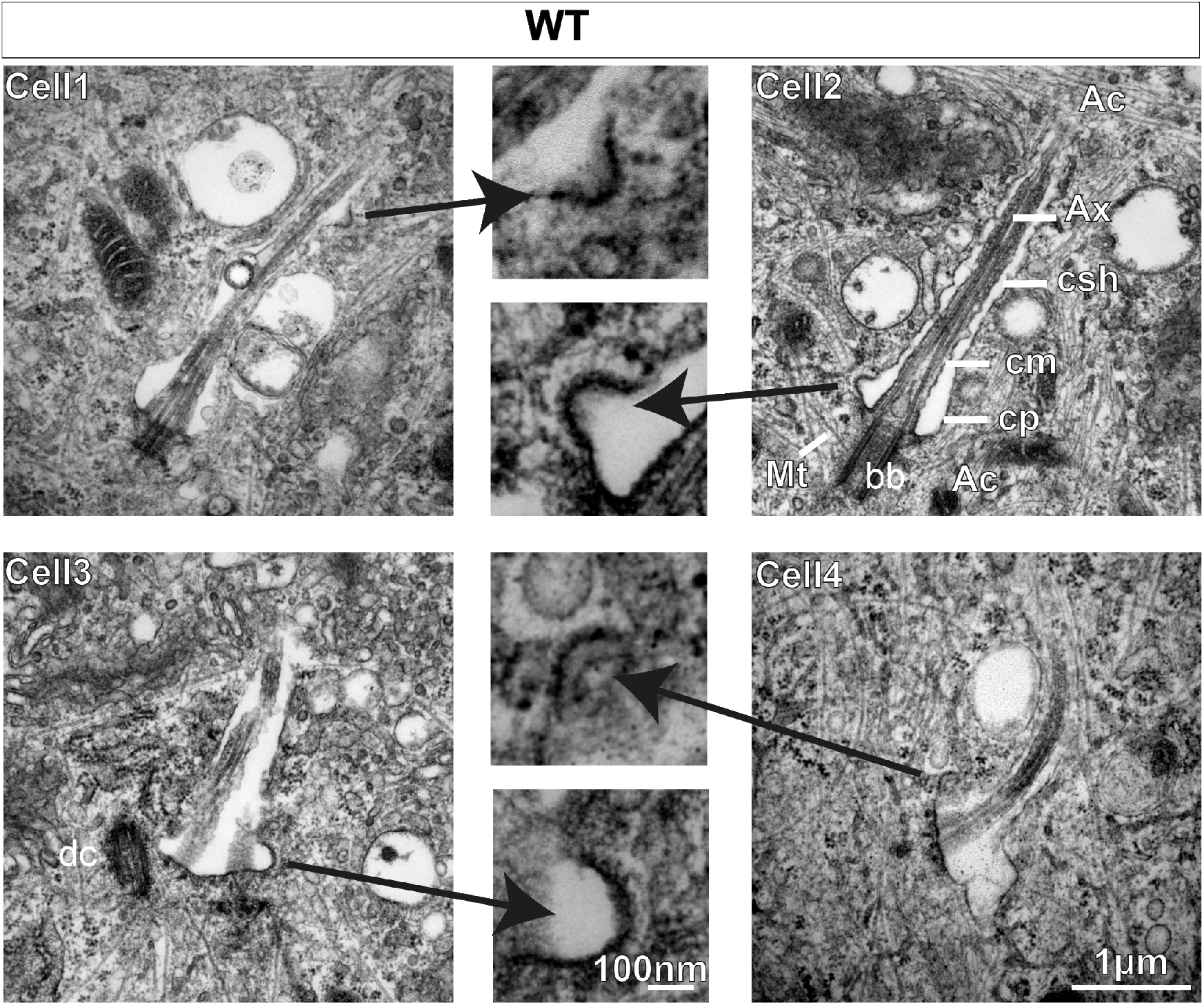
Vesicles with electron-dense coats are observed protruding/fusing with the ciliary sheath in WT MEFs. 24 hr serum-starved WT MEFs are processed for TEM imaging. TEM micrographs of 70 nm sections show vesicles fusing with or protruding from the ciliary sheath, mostly at the ciliary pocket and less along the length. Vesicles are enlarged in the middle panel. Other structures pointed by straight lines are actin filaments (Ac), microtubules (Mt), axoneme (Ax), ciliary sheath (csh), ciliary membrane (cm), ciliary pocket (cp), basal body (bb), daughter centriole (dc). Scale bars = 1 µm in the side lanes and 100 nm in the middle lane.

**Figure 7–Figure supplement 2.**
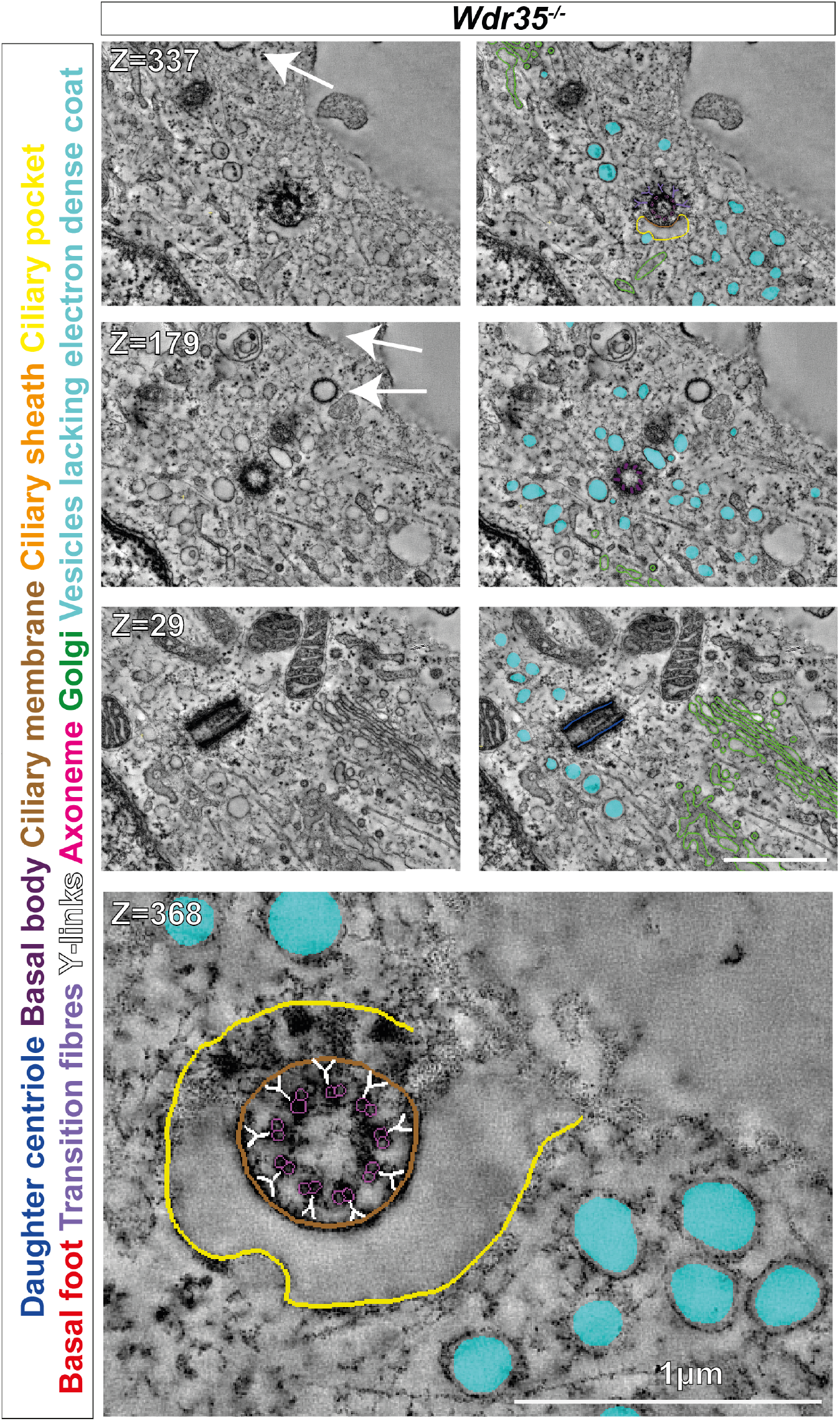
Retrograde dynein motor mutant has a different ciliary structure defect than Wdr35 mutants. After 24 hours of serum starvation, the tilt series was made for 300 nm Wdr35^−/−^TEM samples. Z-stacks from 900 nm serial tomograms is color-coded, highlighting the daughter centriole (dark blue), basal body (purple), ciliary membrane (brown), ciliary sheath (orange), ciliary pocket (yellow), basal foot (red), transition fibres (periwinkle), Y-links (white), axonemal microtubules (magenta), Golgi (green), and vesicles around the cilia (cyan) (Movie 6). Images in the left panel are segmented in the right panel. Uncoated vesicles (cyan) accumulate around mutant cilia but fail to fuse with it. Transition zone (TZ) appeared intact in Wdr35^−/−^mutants. Enlarged TZ in the last panel show no disturbance in (9+0) microtubule doublet arrangement and Y-links connecting axoneme to cilia membrane. The clathrin-coated vesicle can be seen invaginating from the plasma membrane are shown by arrows in the upper two left panels. Scale bars = 1 µm.

**Figure 7–Figure supplement 3.**
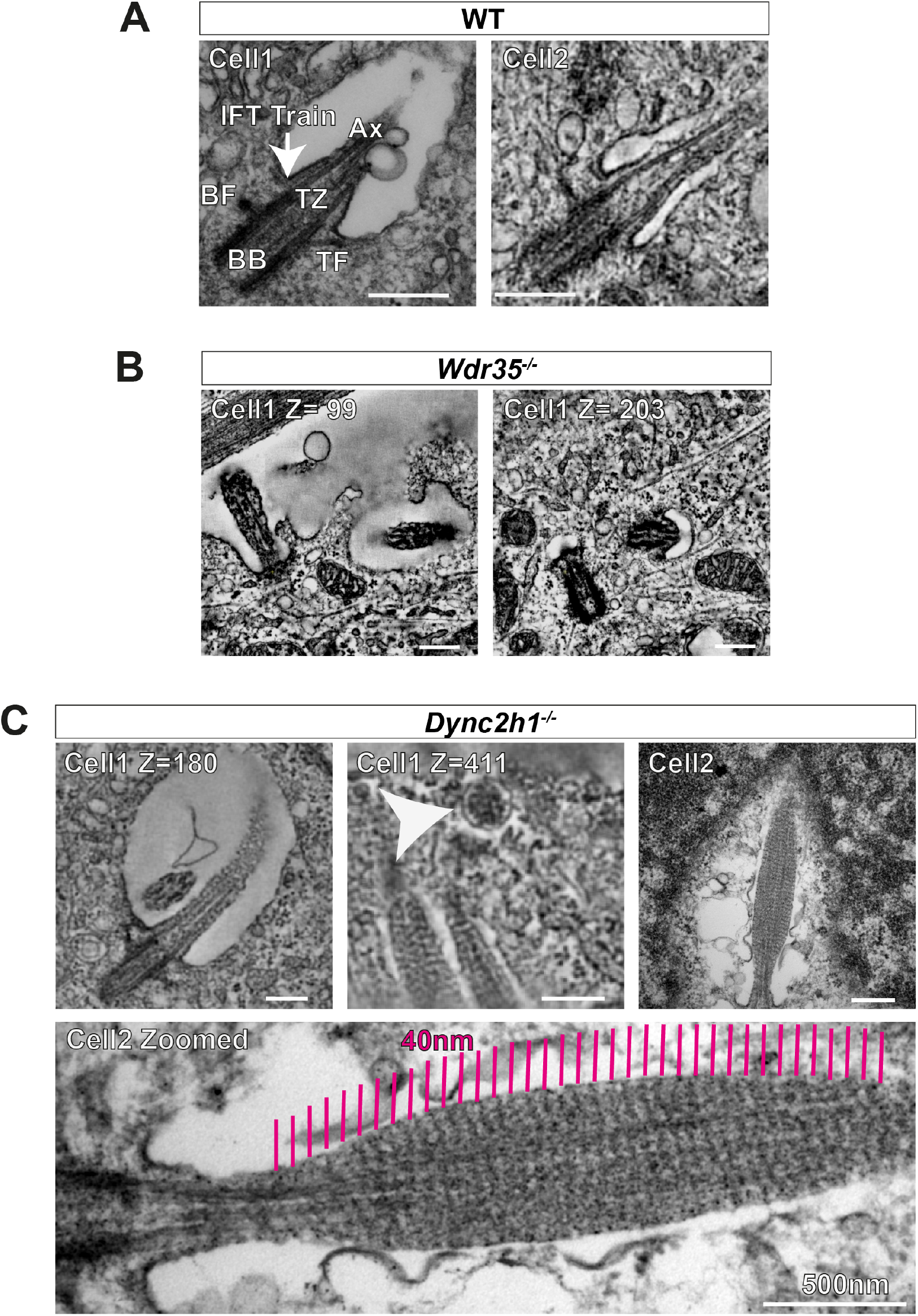
Retrograde dynein motor mutant has a different ciliary structure defect than Wdr35 mutants. (A) 70 nm (cell 1) TEM micrograph and a Z-stack from a tomogram of 300 nm WT MEF showing cilia ultrastructure; basal body (BB), transition zone (TZ), axoneme (Ax), transition fibres (TF), basal foot (BF) and IFT train. The arrow points at the IFT train entering cilia at the ciliary pocket stacked between the axoneme and the ciliary membrane. (B) Z-stack from a serial tomogram reconstructed from 600 nm thick section of Wdr35^−/−^MEFs, the ciliary membrane is less well-defined, and microtubules in the axoneme are disrupted, and periciliary vesicles accumulate around cilia. (C) Z-stack from a serial tomogram of a 900nm thick section (cell 1-Movie 7) and TEM micrograph of 70 nm section (cell 2) of Dync2h1^−/−^MEFs has a striped pattern with a periodicity of 40 nm apparent throughout the length of the cilium. Cell 2 is enlarged to show the same striped pattern (magenta lines). The arrowhead points at the exosome budding from the tip of Dync2h1^−/−^cilium in cell1 (Movie 7). Scale bars= 250 nm, and zoomed out last image is 500 nm.

**Figure 8–Figure supplement 1.**
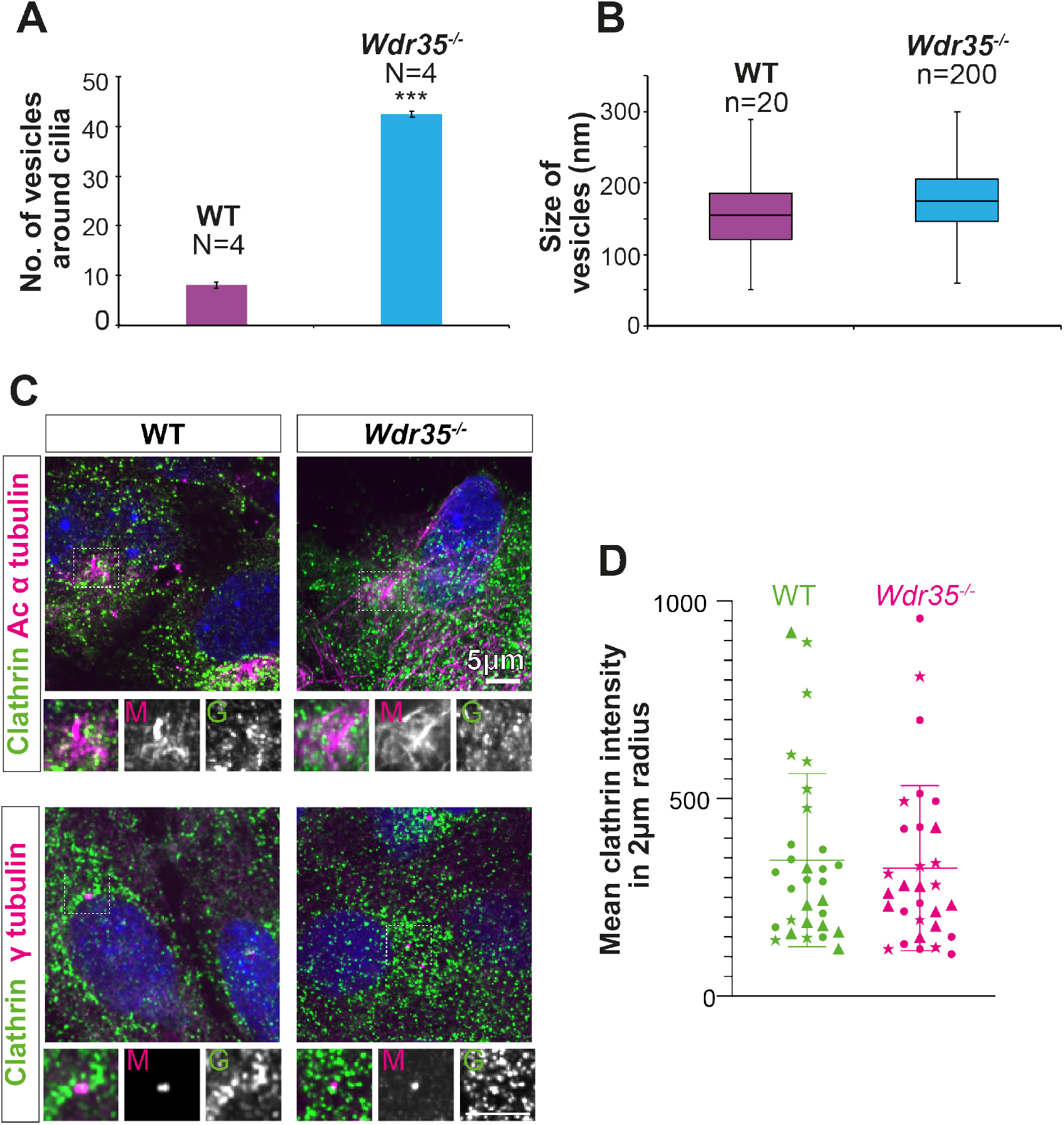
Increased periciliary vesicles in Wdr35 mutant cells are unlikely to be clathrin-based as number and distribution of clathrin-positive foci remains unchanged. (A) The average number of vesicles around cilia in control and Wdr35^−/−^cells, counted in a volume of 2 µm radius around cilia in TEM tomograms show ten times more vesicles in Wdr35^−/−^. N= number of whole-cell volume tomograms per genotype. (B) The size of the periciliary vesicles did not show a significant difference between control and Wdr35^−/−^. n= number of vesicles. The paucity of vesicles around Dync2h1^−/−^cilia prohibited quantification. (C) 24 hr serum-starved cells stained for clathrin antibody (green) and acetylated α tubulin (upper panel) and γtubulin (lower panel) antibodies (magenta) do not show any difference in the distribution of clathrin around cilia. Scale bars = 5 µm. (D) No difference in the mean intensity of clathrin foci quantified in a volume of 2 µm radius around the base of cilia. n= 30 cells (3 biological replicates shown by different shapes each). Asterisk denotes significant p-value from t-test: (*, P<0.05), (**, P<0.001), (***, P<0.0001).

